# Australian giant kelp genome assemblies show distinct Southern Hemisphere genetics

**DOI:** 10.64898/2026.02.20.707121

**Authors:** Hugo J. Scharfenstein, Andrew Carroll, Cintia Iha, Jakop Schwoerbel, Rebecca Jordan, Anusuya Willis

## Abstract

Giant kelp, *Macrocystis pyrifera,* occurs across northern and southern Hemisphere temperate coasts and is at high risk from ocean warming. Few giant kelp forests remain across the Southeast Australian shelf, while a handful of forests are actively being restored. Genomic resources can greatly aid in the conservation of remnant populations and enhance restoration efforts. Reference genomes are a fundamental resource as they are a prerequisite to, or enhance, many analyses used in conservation genomics. A single reference genome is available for giant kelp, assembled from a Californian haploid specimen. However, increasing evidence of genetic divergence between Northern and Southern Hemisphere populations highlights the need for regionally representative reference genomes. Here, we present two genome assemblies from the diploid vegetative tissue of Australian giant kelp specimens. We performed *de novo* genome assembly using long-read sequencing (PacBio HiFi and ONT R10.4 Simplex) and scaffolded the assemblies with the ONT reads, assembling 98-99% of the genomes into 35 pseudo-chromosomes. Genome sizes ranged from 528-534 Mbp, with BUSCO completeness scores of 97-98% and QV scores of 51-52. Genome annotation identified 17,565-17,800 genes in the Australian genomes. Genomic divergence between the Australian and Californian giant kelp genomes was seven-fold greater than between Australian genomes (1.5% vs 0.2%), supporting a Northern–Southern Hemisphere genetic divergence. Functional divergence was also observed between Australian and Californian genomes, reflected by differing patterns of enrichment in gene ontologies linked to energy metabolism, proteostasis and stress responses. These two new genome assemblies will serve as valuable resources for ongoing research into Southern Hemisphere giant kelp genetics, while providing the basis for genomic-guided conservation and restoration of remnant giant kelp forests in Australia.

## Background

Kelp are large brown seaweeds that are key ecosystem engineers of coastal temperate reefs. They form underwater forests that provide fundamental ecosystem services (Eger et al., 2023), are harvested and cultured for food and other products (Forbes et al., 2025) and are anchored in cultural practices (Thurstan et al., 2018). *Macrocystis pyrifera,* also known as giant kelp, is the largest kelp species that can form towering structures over 30m high. *M. pyrifera* is broadly distributed, spanning the North and South American pacific coasts, New Zealand, Southeast Australia and the subantarctic islands (Gonzalez-Aragon et al., 2024).

In Australia, *M. pyrifera* is found along the coasts of the states of South Australia, Victoria and Tasmania. In the global geographical context, Australian and New Zealand populations are the furthest removed from the ancestral populations in the northeast pacific. Over the last decades, giant kelp coverage has drastically declined throughout its Southeast Australian range (Butler et al., 2020). This population collapse is particularly evident in East Tasmania, an ocean warming hotspot (Oliver & Holbrook, 2014), where *M. pyrifera* cover has decreased by 95-98% since 1940 (C. R. Johnson et al., 2011). This decline is primarily attributed to the southward extension of the East Australian Current, bringing warmer, nutrient poor waters to East Tasmania, especially to Northeast Tasmania (Oliver & Holbrook, 2014). This has exacerbated the frequency and severity of marine heatwaves in the area (Oliver et al., 2019) and enabled the range expansion of the long-spined sea urchin *Centrostephanus rodgersii*, which prevents giant kelp recruitment and causes urchin barrens (Ling, 2008). To reverse this drastic decline of this key habitat forming species, extensive restoration of giant kelp forests is needed. Thankfully, [such efforts](https://www.utas.edu.au/about/news-and-stories/articles/2024/working-together-to-safeguard-threatened-giant-kelp-forests) are now underway along East Tasmania and beyond, such as on the Pacific coast of North America.

Genomic resources are increasingly incorporated into the conservation of threatened species, providing valuable information for their management to ensure preservation of genetic diversity or population structure (Dussex et al., 2021; Morin et al., 2021; Wood et al., 2020). The decreasing cost of high-throughput sequencing and democratisation of analysis software have increased the availability of genome-scale datasets, providing genomic perspectives of the genetic diversity and historical demography of threatened populations/species (W. E. Johnson et al., 2014). A reference genome (i.e., a highly contiguous, accurate, and annotated genome assembly) is fundamental to maximising use of high-throughput sequencing data, enabling genome mapping for subsequent genetics analyses. Additionally, high-quality reference genomes can aid in the development of molecular markers, which provide a rapid and cost-effective tool to screen individuals for traits of interest (Brandies et al., 2019).

A single reference genome of *M. pyrifera* has been assembled from a haploid Californian specimen (Diesel et al., 2023). A recent population genomics study of giant kelp was able to leverage this genome with high-throughput sequencing data, revealing multiple genetic health risks in populations from western Canada and USA (Bemmels et al., 2025), demonstrating the value of a reference genome and genetic data. Developing such genomic resources in Tasmania is urgently needed to inform the conservation of remnant giant kelp populations and enhance local restoration efforts.

No reference genome is currently available for giant kelp from the Southern Hemisphere, let alone Australia, which limits population genetics studies to rely on the Californian genome. Additionally, recent studies investigating the genetics of *M. pyrifera* have re-ignited the taxonomic debate surrounding giant kelp, with significant genetic divergence repeatedly reported between Northern and Southern hemisphere populations (Assis et al., 2023; Bemmels et al., 2025; Gonzalez et al., 2023). Given the apparent genetic divergence between Northern and Southern hemisphere giant kelp, it is vitally important to use a Southern Hemisphere reference genome for genetics analysis, or risk producing misleading information (Formenti et al., 2022). Creating a reference genome for Australia is also essential to inform local conservation and restoration efforts. Further reinforcing the need for a local reference genome is the notion that no single genome can account for all the genetic diversity within a species (Liao et al., 2023). The availability of multiple reference genomes for a species can reduce errors in genomic characterisation (Valiente-Mullor et al., 2021), while providing the building blocks of a species pangenome.

To address this knowledge gap, we assembled the first giant kelp genomes from the Southern Hemisphere to provide a valuable resource to reduce individual bias for future giant kelp genomics studies. We conducted long-read sequencing (PacBio HiFi and ONT R10.4 Simplex) to assemble the genomes of two Australian giant kelp individuals to quasi chromosomal-level. We compared genome structure and function between these two Australian genomes and with the available reference from California.

## Methods

### Sample collection

Two giant kelp individuals from the Southeast Australian Shelf were sampled in 2024, one at Crayfish Bay, Victoria (sample ID: OTW_2, latitude: –38.8548, longitude: 143.5376) and the other around 600km Southeast at Shelly Point, Tasmania (SHL_1, –41.4353, 148.2785), under scientific research permit n°23090 (Department of Natural Resources and Environment Tasmania). Hereafter, the samples will be referred to by their region of origin (i.e., Victoria and Tasmania). For each individual, vegetative tissue from the apical blades were sampled, wiped with a 10% (v/v) bleach solution, rinsed under freshwater and blotted dry. The cleaned apical blades were snap frozen in liquid nitrogen and stored at –80°C within 24 hours of sampling.

### DNA extractions

High molecular weight DNA was extracted following a protocol modified from Diesel et al. (2023) (see Supplementary Information for the detailed DNA extraction protocol). Briefly, the flash frozen giant kelp tissue was ground to fine powder in a mortar topped up with liquid nitrogen. The powder was transferred to a CTAB lysis buffer, modified from Doyle & Doyle’s (1987) original recipe, composed of: 1.2 M NaCl (CAS: 7647-14-5), 20 mM EDTA (CAS: 60-00-4), 100 mM Tris-HCl (CAS: 1185-53-1), 2% (w/v) CTAB (CAS: 57-09-0), 2% (w/v) PVP-40 (CAS: 9003-39-8), 1% (w/v) PEG 8,000 (CAS: 25322-68-3), 1% (v/v) β-mercaptoethanol (CAS: 60-24-2) and 100 µg/ml Proteinase K (EO0492; Thermo Fisher Scientific). The ground tissue suspended in CTAB lysis buffer was incubated for 1 hr at 55°C, inverting every 15 min.

The lysate was centrifuged for 5 min and the supernatant mixed with an equal volume of Chloroform: Isoamyl alcohol 24:1 (C0549, Sigma-Aldrich). The aqueous and organic phases were separated through centrifugation at 4°C for 15 min. The upper aqueous phase was incubated for 30 min at 37°C with RNAse A (12091021; Invitrogen) at a final concentration of 80 µg/ml. The lysate was mixed a second time with Chloroform: Isoamyl alcohol and centrifuged at 4°C for 15 min. The upper aqueous phase was mixed with 0.4x volume of 5 M Potassium Acetate (CAS: 127-08-2) and incubated for 20 min on ice. The lysate was centrifuged at 4°C for 45 min. The resulting supernatant was mixed with 0.1x volume of 3 M Sodium Acetate (CAS: 127-09-3) and 0.8x volume of Isopropanol (CAS: 67-63-0). After a 30 min incubation at room temperature, the DNA was precipitated out of solution by centrifugation for 30 min. The pellet was washed twice with 70% Ethanol, air-dried for 3-5 min and resuspended in TE buffer.

### Sequencing

Two rounds of PacBio sequencing were carried out to overcome the relatively low output obtained from a single sequencing run. See Table S1 for a detailed breakdown of DNA input and sequencing output for each sample/run/sequence type. The samples were sequenced on the same PacBio Revio platform at the Biomolecular Resource Facility (Australian National University), Canberra, Australia for both runs. One round of ONT sequencing was carried out on a PromethION platform at the Biomolecular Resource Facility. For each sample and sequencing run, 3.5-11.9 µg of DNA was sent for library preparation, which was carried out at the sequencing facility. Genomic DNA size was validated by Femto Pulse (Agilent), with a median fragment length of 25-50 Kbp (Table S1). Size selection was performed via Blue Pippin (Sage Science), with DNA recovery rates varying from 14-33% (Table S1). Samples were multiplexed on either one SMRT flow cell, for PacBio sequencing, or on one R10.4.1 flow cell, for ONT sequencing.

Both samples were sequenced on an Illumina NovaSeq 6000 platform (2×150 bp) by Azenta Life Sciences. For each sample, three replicate libraries were prepared. For each library, 200 ng of genomic DNA underwent library preparation at the sequencing facility using a VAHTS Universal Plus DNA Library Prep Kit for Illumina V2 (Vazyme), according to the manufacturer’s protocol. Each library was sequenced on a separate Illumina NovaSeq 6000 lane, with each sample multiplexed on the same lane.

### Sequencing read processing

HiFi reads were screened for contaminants using *fcs-gx* v0.5.5 (Astashyn et al., 2024) with the ‘--mask-transposons’ option turned off. HiFi reads were removed if they were identified as ‘EXCLUDE’ by *fcs-gx* (Table S2). Valid ONT R10.4 reads (i.e., minimum Phred quality score of 10) were discarded if they were shorter than 10 Kbp (Table S2).

### Organellar genome assembly

Mitochondria and chloroplast genomes were assembled for each sample from the filtered HiFi reads using *oatk* v1.0 (Zhou et al., 2024). The resulting organellar genomes were annotated using *GeSeq* (Tillich et al., 2017) and visualised with *OGDRAW* (Greiner et al., 2019).

### Nuclear genome assembly

A summary of the software and settings used in the nuclear genome assembly and annotation pipelines is provided in Table S3, while an overview of the workflow is provided in Fig. S1. Primary and alternate assemblies were generated for both samples using *hifiasm* v0.25.0 (Cheng et al., 2021) with filtered HiFi reads, while using ONT reads as input for scaffolding (using the –-ul option).

The nuclear assemblies were then screened for the presence of organellar contigs. A BLAST search was carried out against a custom database composed of giant kelp organellar genomes assembled from the HiFi reads, as well as giant kelp organellar genomes retrieved from Iha et al. (2023), Starko et al. (2021), Chen et al., (2019) and GenBank (Accession ID: MW899036.1). In addition, the filtered HiFi reads were mapped back to the primary and alternate assemblies using *minimap2* v2.29 (Li, 2018) to determine the coverage of each contig. A contig was removed if it presented a high similarity to the organellar genome database (i.e., an e-value lower than 1e-10 and qcov greater than 60%) and a high coverage (i.e., 3x greater than the sequencing coverage) or a GC content below 0.4. The assemblies were then screened for contaminants using *blobtoolkit* v4.2.1 (Laetsch & Blaxter, 2017). For each assembly, we obtained results from searches against the BLAST nucleotide database (v5, March 2025 release), UniProt Reference Proteomes database (April 2025 release) and BUSCO database (v5, March 2025 release). The results were added to *blobtoolkit* and contigs were filtered based on GC content (contigs with a GC content between 0.4–0.6 were kept) and taxonomy (contigs were kept if they were classified as Phaeophyceae, Bigyra or no-hit). See Fig. S2-S3 for blobplots.

The filtered assemblies underwent further duplicate purging using *purge_dups* v1.2.5 (Guan et al., 2020), with cutoffs for coverage thresholds set manually (Table S3). The purged primary assemblies were scaffolded with ONT reads using *ntLink* v1.3.11 (Coombe et al., 2023) in sensitive mode and with the minimum number of anchoring reads doubled (a=2) for more stringent scaffolding. Gaps in the scaffolded assemblies were filled with *TGS-GapCloser* v1.2.1 (Xu et al., 2020) using ONT reads corrected with Illumina reads by *pilon* v1.24 (B. J. Walker et al., 2014). All assemblies underwent a final screening for contaminants using *fcs-gx* v0.5.5 before submission to NCBI’s Genomes database (see Data Availability section for further detail).

### Genome annotation

The primary assemblies were soft masked sequentially using *RepeatMasker* v4.1.9 (Tarailo-Graovac & Chen, 2009). Firstly, with Dfam’s database (release 3.9; Hubley et al., 2016), specifying Phaeophyceae for taxa search, and secondly with repeats identified by *RepeatModeler* v2.0.2a (Flynn et al., 2020). Prior to the second round of masking, gene fragments were removed from the repeat library generated by *RepeatModeler.* This was done by removing transposons from two clade-partitioned (Stramenopiles and Viridiplantae) OrthoDB v12 (Kuznetsov et al., 2023) protein datasets using *TransposonPSI* v1.0.0 (Riehl et al., 2022). A BLAST search was conducted between the repeats identified by *RepeatModeler* and the transposon-free protein datasets, removing hits with *ProtExcluder* (Shu, 2019).

Prediction of protein coding gene structures was carried out using *BRAKER* v3.0.7 (Hoff et al., 2016), following the BRAKER3 pipeline. As input for transcriptomic evidence, we retrieved 24 RNAseq giant kelp datasets from NCBI’S Sequence Read Archive (BioProject accession IDs: PRJNA353611, PRJNA322132, PRJNA1116209). As input for protein evidence, we combined the OrthoDB v12 dataset with protein datasets from JGI’s Genome Portal for four Laminariales species: *M. pyrifera* (Diesel et al., 2023, Portal ID: Macrocystis pyrifera CI_03 v1.0), *Saccharina japonica* (Ye et al., 2015, Portal ID: Saccharina japonica str. Ja), *Undaria pinnatifida* (Shan et al., 2020, Portal ID: Undaria pinnatifida M23) and *Nereocystis luetkeana* (Alves-Lima et al., 2025, Portal ID: Nereocystis luetkeana NB12FB3 v1.0). Transcriptomic and protein datasets then served as extrinsic evidence for training of *GeneMark-ETP* v1.02 (Brůna et al., 2024) and *AUGUSTUS* v3.4.0, which also incorporated the output from *GeneMark-ETP*. The final gene set was predicted by *TSEBRA* v1.1.2.5 (Gabriel et al., 2021) with the combined results from *GeneMark-ETP* and *AUGUSTUS*. Functional annotation of the gene sets was performed using all available analyses from *InterProScan* v5.72-103.0 (Jones et al., 2014), as well as Phobius1.01 (Käll et al., 2004), SignalP v4.1 (Petersen et al., 2011) and THMM v2.0 (Krogh et al., 2001).

Gene prediction and functional annotation were also performed on the available masked Californian reference genome to provide a like-for-like comparison of gene content with the Australian genomes for orthology and gene ontology analyses.

### Assembly quality and genome features

Assembly contiguity was assessed via *assembly-stats* v1.0.1. Gene content completeness was evaluated via *BUSCO* (v5.8.0; Manni et al., 2021) against the stramenopiles_odb10 (2024-01-08) dataset, using Augustus v3.5.0 (Stanke et al., 2006) as gene predictor and with the flag ‘--long’. QV scores of the final assemblies were assessed with *yak* v0.1 (https://github.com/lh3/yak).

*Circos* plots (v0.69.9; (Krzywinski et al., 2009) of the primary assemblies were generated to depict: scaffold size, average GC content over 100 Kbp windows (using *bedtools* v2.31.1; Quinlan & Hall, 2010), read coverage over 10 Kbp bins (generated with *minimap2* and *bedtools*), number of interspersed repeats greater than 10 Kbp (previously identified by *RepeatMasker*) over 10 Kbp bins and segmental duplications above 15 Kbp between contigs (identified with *asgart* v2.4.0; Delehelle et al., 2018).

### Comparative genomics

Divergence between the Victorian, Tasmanian and Californian genomes was estimated using *Mash* v2.3 (Ondov et al., 2016). We then performed a synteny analysis between all three genomes using *ntSynt* v1.0.2 (Coombe et al., 2024), which was visualised using *ntSynt-viz* v1.0 (Coombe et al., 2025). Using *OrthoFinder* v3.0.1b1 (Emms et al., 2025), we compared orthology between all three *M. pyrifera* genomes (Victorian, Tasmanian and California) and three other Laminariales: *Saccharina japonica* (Ye et al., 2015), *Undaria pinnatifida* (Shan et al., 2020) and *Nereocystis luetkeana* (Alves-Lima et al., 2025).

An enrichment analysis was carried out with the gene ontology (GO) terms previously identified by *InterProScan*. Prior to enrichment analysis, GO annotations were collapsed to the gene level by merging all gene isoform associated GO terms and removing duplicate GO terms. A gene set of interest was prepared for each assembly by selecting GO terms present in greater abundance in the focal assembly than in a second assembly. This process was repeated for each possible pairwise comparison between assemblies. Gene universes were prepared for each pairwise comparison by combining gene sets with GO annotations from both assemblies. GO enrichment analysis was then performed with *topGo* v2.58.0 (Alexa & Rahnenführer, 2025), using the algorithm *weight01* and Fisher’s exact test to identify significantly enriched GO terms in each pairwise comparison. To focus on broader trends, a ‘nodeSize’ of 10 was used to prune the GO hierarchy from terms with less than 10 annotated genes.

## Results

### Sequencing

PacBio Revio sequencing generated 3.5 M (44.7 Gbp) and 5 M (62.5 Gbp) HiFi reads for the Victorian and Tasmanian samples, respectively (Table S1). This corresponds to an 83x and 116x coverage of the 537 Mbp haploid *M. pyrifera* genome assembled by Diesel et al. (2023). A N50 of 14.4 Kbp was obtained across both samples and runs (Table S1). Contaminant filtering led to the removal of 64,604 (812 Mbp) and 4,980 (57 Mbp) HiFi reads from the Victorian and Tasmanian datasets.

ONT sequencing generated 0.55 M (12.15 Gbp) and 0.6 M (14.5 Gbp) R10.4 reads with a minimum Q score of 10 for Victorian and Tasmanian samples, respectively (Table S1). A N50 of 31 Kbp was obtained across both samples. Length filtering led to the removal of 0.16 M (0.68 Gbp) and 0.11 M (0.50 Gbp) ONT reads (Table S2).

### Organellar genomes

The mitochondrial genomes of the Victorian and Tasmanian samples are 37,313 and 37,297 bp long, respectively, with a GC content of 31.82% and 31.87% (Fig. S4). Both mitochondrial genomes contain 3 rRNA-coding genes (LSU rRNA, SSU rRNA, 5S rRNA), but differ in their number of distinct tRNA loci (25 for the Victorian genome vs 22 for the Tasmanian genome) and unique protein-coding genes (31 for the Victorian genome vs 27 for the Tasmanian genome). See Table S4 for a list of gene features in each mitochondrial genome.

The chloroplast genomes are 130,184 and 130,212 bp long for the Victorian and Tasmanian samples, respectively. GC content is at 30.88% for both samples (Fig. S5). The large single-copy region is 78,069 bp long for the Victorian genome, compared to 53,755 bp long for the Tasmanian genome, while the small single-copy region is 42,875 bp long for the Victorian genome (42,903 bp for the Tasmanian genome). Both chloroplast genomes share 3 rRNA-coding genes, 25 tRNA-coding genes (total of 26 tRNAs for each genome) and 50 protein-coding genes (total of 57 and 61 genes for the Victorian genome and Tasmanian genome, respectively) (Table S5).

### Nuclear genome assembly

*De novo* assembly generated primary assemblies at 557 Mbp (311 contigs) for the Victorian sample and 557 Mbp (549 contigs) for the Tasmanian sample, with corresponding N50 values of 7.1 Mbp and 7.4 Mbp (see Table S6 for assembly statistics throughout the assembly pipeline). The Tasmanian alternate assembly was three times the size (1.44 Gbp) of the Victorian alternate assembly (0.48 Gbp). Organellar contig and contaminant filtering led to the removal of 175 contigs (9.7 Mbp) and 330 contigs (19.6 Mbp) from the Victorian and Tasmanian primary assemblies, respectively (Table S6). Scaffolding resulted in primary assemblies with 39 scaffolds/contigs (30 scaffolds, 9 contigs) and 49 scaffolds/contigs (23 scaffolds, 26 contigs) for the Victorian and Tasmanian genomes, respectively (Table 1). 99.3 and 97.5% of the Victorian and Tasmanian genomes assembled into 35 pseudo-chromosomes, respectively. Final genome sizes of 534 and 528 Mbp were obtained for Victorian and Tasmanian primary assemblies.

**Table 1:**
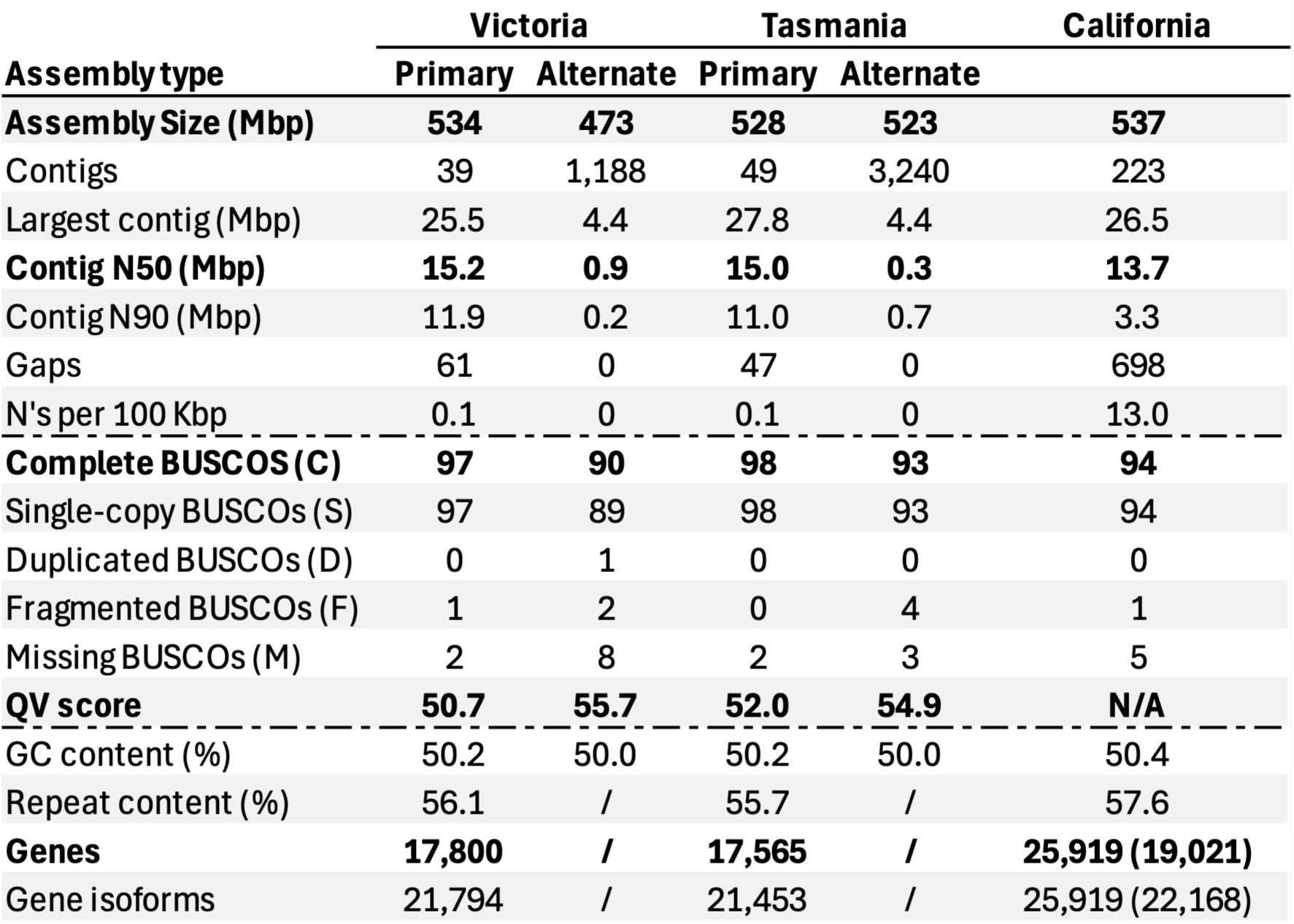
Summary statistics of the Victorian and Tasmanian giant kelp genome assemblies from this study and the Californian reference genome from Diesel *et al*. (2023). For the Californian assembly, metrics in parentheses were estimated in this study.

Gene content completeness, assessed via BUSCO analysis, was estimated at 97% and 98% for the Victorian and Tasmanian primary assemblies, respectively (Table 1). The N50 of both primary assemblies was assessed around 15 Mbp, with 47-61 gaps recorded. QV scores of 51 and 52 were obtained for the Victorian and Tasmanian primary assemblies, respectively.

### Genome features

GC content was estimated at 50.20% and 50.22% for the Victorian and Tasmanian assemblies, respectively, compared to 50.37% for the Californian reference (Table 1). Repetitive elements made up 56.1% and 55.7% of the Victorian and Tasmanian genomes, respectively, compared to 57.6% reported for the Californian reference (Diesel et al., 2023). Interspersed repeats represented the near totality (87-88%) of repetitive elements identified in the Australian genomes (Table S7). Gene prediction identified 17,800 genes (21,794 gene isoforms) and 17,565 genes (21,453 gene isoforms) for the Victorian and Tasmanian genomes, respectively. In comparison, 19,021 genes (22,168 gene isoforms) were predicted for the Californian genome.

### Genome comparisons

Genome divergence between the Victorian and Tasmanian assemblies was estimated by *Mash* at 0.2% (912/1,000 matching hashes). Both Australian genomes displayed a 1.5% divergence with the Californian reference genome (568/1,000 matching hashes). In total, 94% (500 Mbp) of the Victorian genome was found to be in synteny with the Tasmanian genome, forming 1,122 synteny blocks with a N50 of 2.2 Mbp. In comparison, 86-87% (461 Mbp) of the Australian and Californian genome structures were conserved.

All three *M. pyrifera* assemblies displayed substantial genome-wide collinearity across their 35 largest scaffolds/contigs (Fig. 3). Rearrangements were observed between Australian genomes, notably at scaffold 20 of the Victorian assembly and scaffolds 8 and 28 of the Tasmanian assembly. Scaffold 2 of the Victorian assembly was split in two in the Tasmanian assembly, though not in the Californian reference. Other structural differences between the Australian and Californian genomes include the splitting of scaffolds 7 and 12 of the Victorian assembly into two in the California assembly.

**Figure 1:**
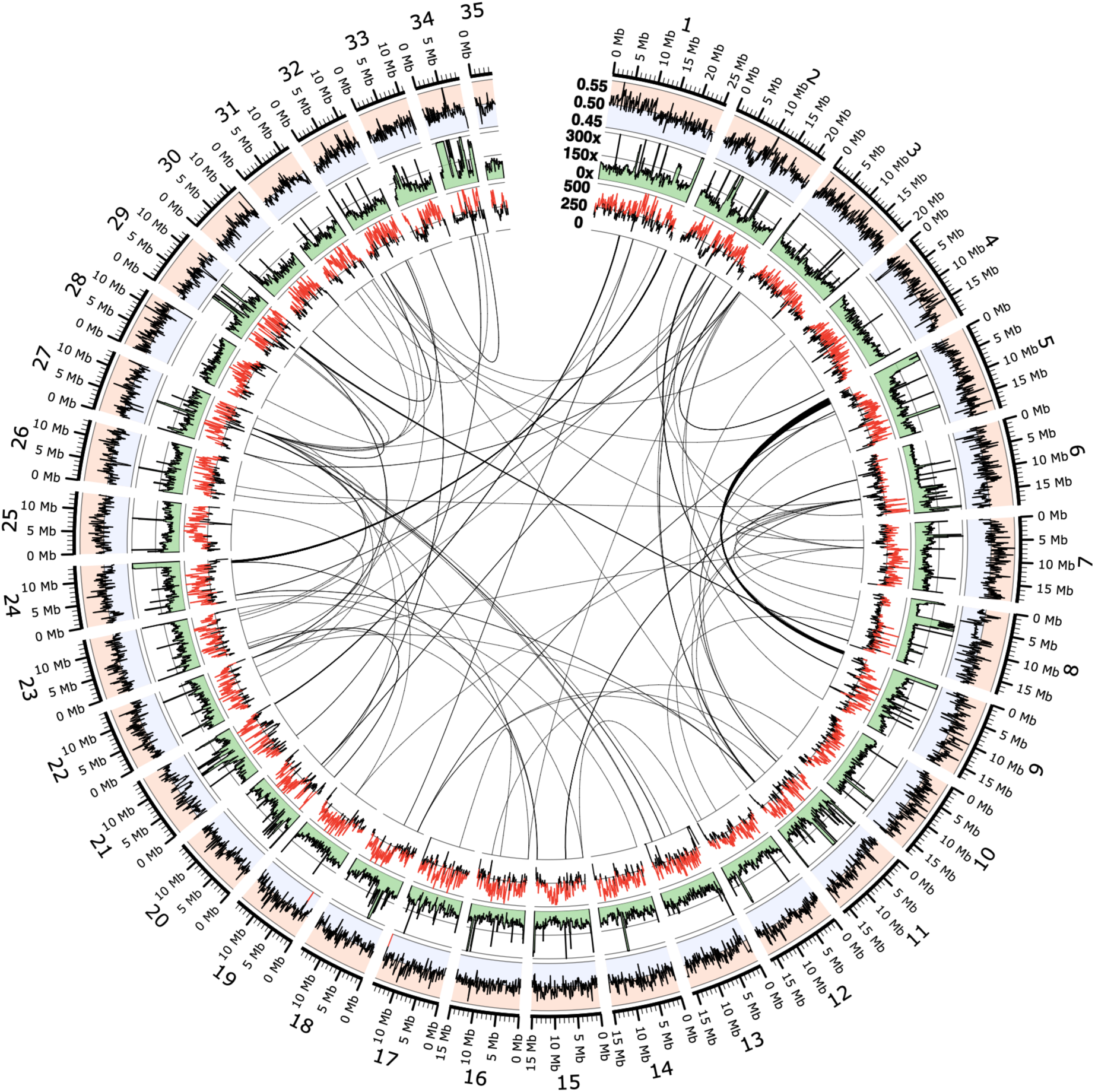
Circos plot of the Victorian primary assembly. Characteristics of the 35 largest scaffolds (equivalent to 99.3% of the assembly) are shown in each concentric circle. Starting from the outermost circle, they show scaffold size in Mbp, GC content over 100 Kbp sliding windows (range: 0.45-0.55), read coverage over 10 Kbp sliding windows (range: 0x-300x), number of interspersed repeats greater than 10 Kbp in size over 10 Kbp windows (range: 0-500) and segmental duplications between scaffolds greater than 15 Kbp.

**Figure 2:**
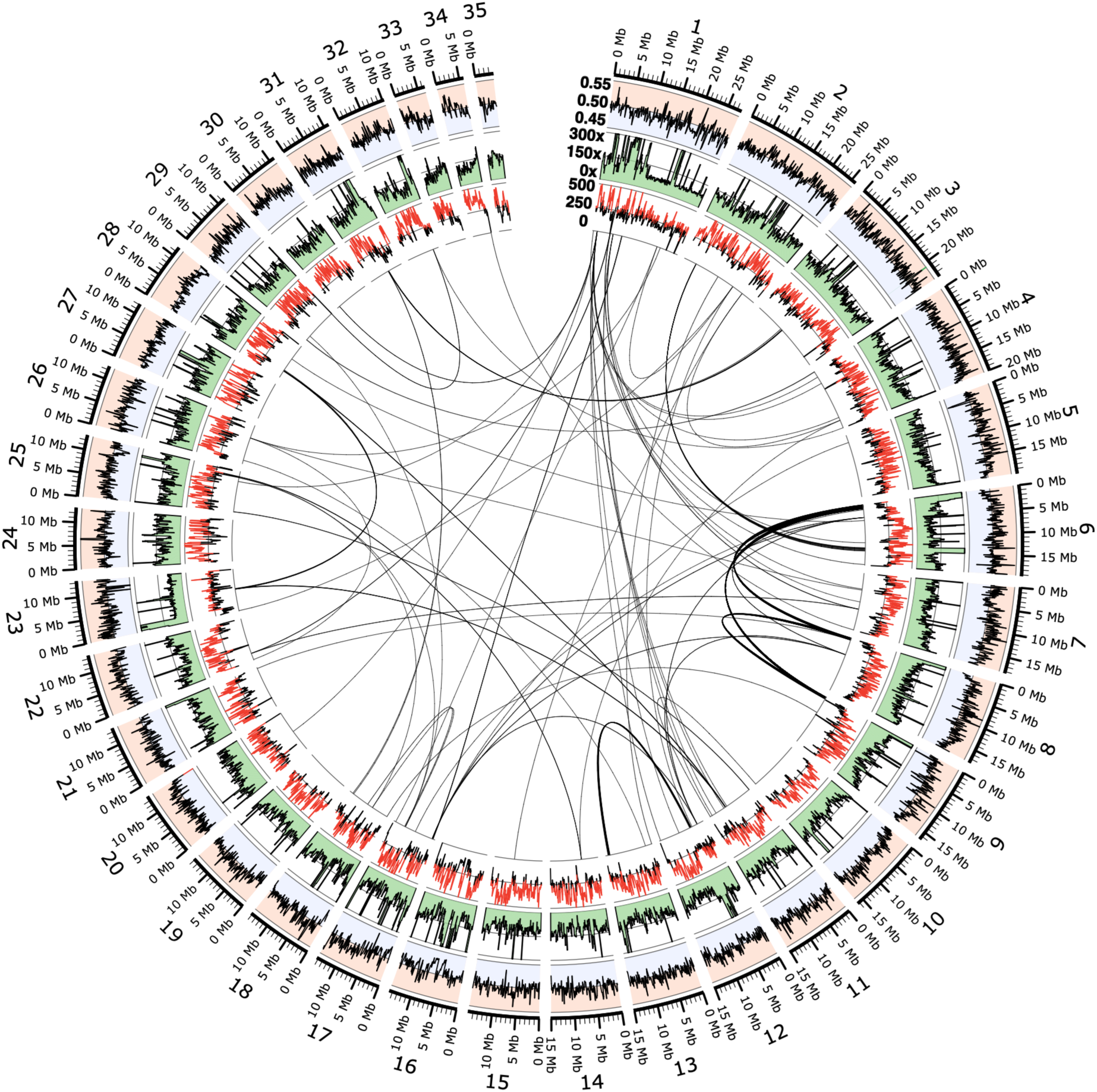
Circos plot of the Tasmanian primary assembly. Characteristics of the 35 largest scaffolds greater (equivalent to 97.5% of the assembly) are shown in each concentric circle. Starting from the outermost circle, they show scaffold size in Mbp, GC content over 100 Kbp sliding windows (range: 0.45-0.55), read coverage over 10 Kbp sliding windows (range: 0x-300x), number of interspersed repeats greater than 10 Kbp in size over 10 Kbp windows (range: 0-500) and segmental duplications between scaffolds greater than 15 Kbp.

**Figure 3:**
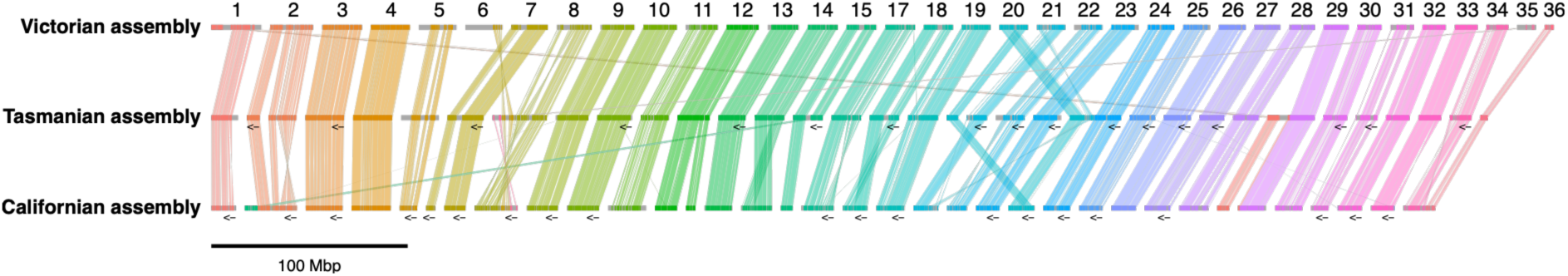
Cross-genome synteny between Australian and Californian genomes. Synteny blocks greater than 5 Mbp are depicted. For each genome, the 35/36 largest scaffolds are shown. Arrows indicate reverse complementation of a synteny block. Each scaffold is represented by a different colour.

Orthology analysis identified 9,703 orthologous groups common to all three giant kelp genomes (Fig. 4A). Five to six times more orthologous groups were shared between the Australian assemblies (2,142) than with the Californian assembly (362-420). Surprisingly, more orthologous groups were shared between the Australian giant kelp genomes and the Laminariales species *N. luetkeana* (10,771-10,822) than with the Californian genome (10,481-10,534) (Fig. 4B). All three giant kelp genomes shared the lowest number of orthologous groups with the *U. pinnatifida* genome.

**Figure 4:**
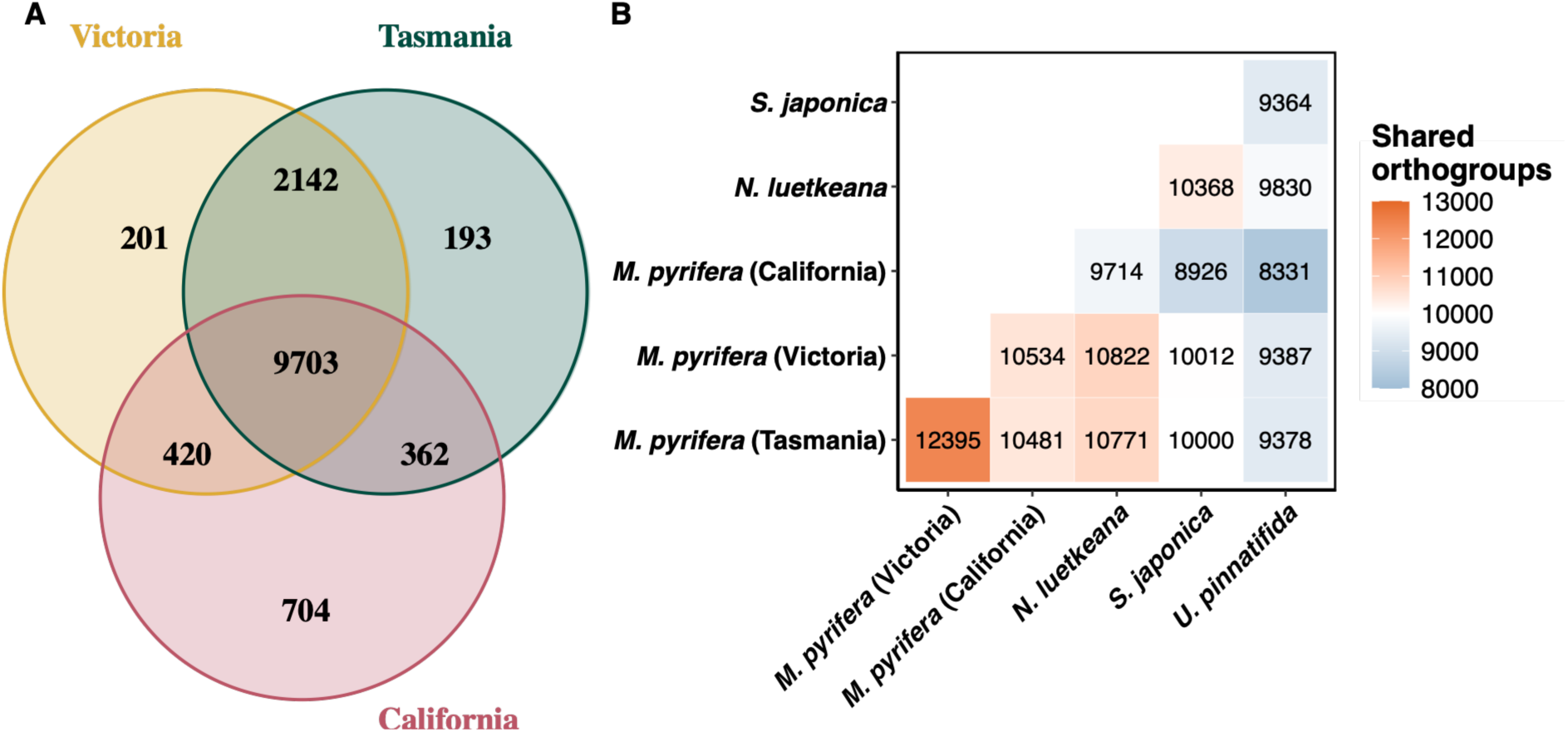
Orthology analysis between genome assemblies from four Laminariales species. **(A)** Venn diagram of shared orthologous groups between all three giant kelp genomes (Victorian, Tasmanian and Californian assemblies). **(B)** Shared orthologous groups between all three giant kelp genomes (*Macrocystis pyrifera*) and three other Laminariales species (*Nereocystis luetkeana*, *Saccharina japonica* and *Undaria pinnatifida*).

Relative to the counterpart Australian assembly, 56 gene ontology (GO) terms were significantly enriched (p_Fisher_<0.01) in the Victorian genome, compared to 33 GO terms in the Tasmanian genome (Table S8). Fewer GO terms (n=13) were significantly enriched in the Australian genomes relative to the Californian genome than conversely (n=114). Compared to the Tasmanian assembly, the Victorian genome showed a distinct enrichment in biological processes related to primary metabolism (glycolytic process and lipid metabolic processes – GO:0006096, GO:0006629) (Fig. 5, Fig. S6). In contrast, the Tasmanian genome was enriched in proteostasis pathways (protein folding, proteolysis, protein ubiquitination – GO:0006457, GO:0006508, GO:0016567), relative to the Victorian genome. The Californian assembly was characterised by an enrichment in oxidative stress response processes (glutathione metabolic process, cell redox homeostasis, cellular response to oxidative stress – GO:0006749, GO:0045454, GO:0034599).

**Figure 5:**
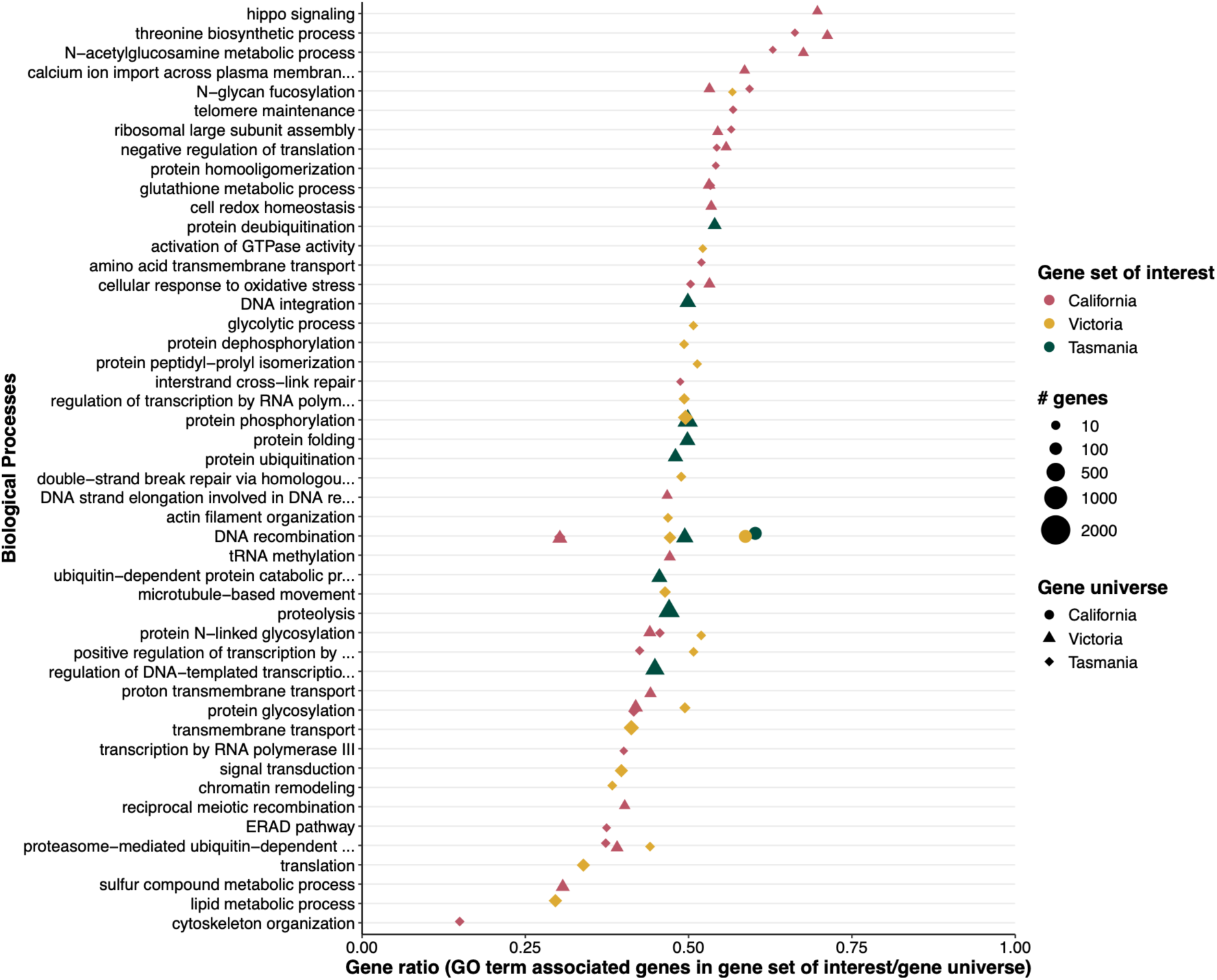
Comparative analysis of gene ontologies (GOs) enriched in Victorian, Tasmanian and Californian giant kelp genomes. For each pairwise comparison, the top 20 significantly enriched GO terms (p_Fisher_<0.01) from the category biological processes are shown here. GO terms pertaining to molecular functions and cellular components can be found in Fig. S6. A list of GO IDs can be found in Table S8. Significant differences were calculated with Fisher’s exact test.

## Discussion

We present the first genomes assembled from Southern Hemisphere *M. pyrifera* specimens using long read sequencing. Both assemblies are highly contiguous, near chromosomal level, and expand upon the current reference genome from California. They also represent the first genomes assembled from the sporophyte (diploid) life stage of giant kelp.

Crucially, these two genomes can serve as the basis for future genetics studies of giant kelp in Australia, and the Southern Hemisphere.

We assembled over 98% of each Australian genome into 35 pseudo-chromosomes, the same number of chromosome-level scaffolds as the Californian genome after Hi-C scaffolding by Diesel et al. (2023). In contrast, 92% of the Californian genome was assembled into 35 pseudo-chromosomes (Diesel et al., 2023). While the Australian and Californian genomes are highly contiguous, neither of the assemblies appears fully resolved at the chromosome-level. Historically, cytological studies have observed haploid giant kelp with ploidy around 14-18 chromosomes (Cole, 1968; Lewis & Neushul, 1994; F. T. Walker, 1952) or 30-32 chromosomes (Lewis & Neushul, 1994; Yabu & Sanbonsuga, 1987), with the large differences in ploidy attributed to parthenogenesis. Cross genome synteny revealed that two scaffolds in the Australian genomes are split in the Californian reference, while one scaffold in the Californian assembly is split in the Australian genomes. The *Undaria pinnatifida* genome, a closely related Laminariales species, was scaffolded into 30 chromosomes with Hi-C sequencing (Shan et al., 2020), further reinforcing the notion that all three giant kelp assemblies are insufficiently scaffolded to reach complete chromosome-level.

In Tasmanian genome we found the alternate assembly (i.e., the assembly representing the alternate haplotype) to be nearly three times the expected genome size, prior to purging (1.44 Gbp). The nature of this larger assembly is unclear, but could be linked to whole genome duplication or be reflective of the presence of one or several additional haplotype(s). Further research is needed to determine the karyotype of this individual and source population, which may explain the larger assembly.

Most genomic features were comparable between the Australian and Californian assemblies. The Australian genomes are similar in size to the Californian reference genome (528-534 Mbp for the Australian assemblies vs 537 Mbp for the Californian assembly), providing further bioinformatic evidence that the haploid genome size is around 530-540 Mbp for giant kelp. Repetitive elements formed around 56% of the Australian giant kelp genomes, comparable to the Californian genome (57%) and other kelp genomes (e.g., 60% for *Saccharina sculpera*; Guo et al., 2025). Gene content was estimated at 17,565-17,800 genes for the Australian genomes. While considerably lower than the 25,919 genes identified by Diesel et al. (2023), re-annotation of the Californian reference using *BRAKER3* identified 19,021 genes, a more comparable number of genes to the Australian assemblies. Hence, this discrepancy in gene content appears to be methodological, rather than biological. It may be attributed to the *BRAKER3* pipeline used here incorporating both protein and transcriptomic evidence, rather than solely transcriptomic evidence as was the case in Diesel et al. (2023), and differences in gene prediction software used.

Greater genomic divergence was observed between the Australian and Californian assemblies than between the Tasmanian and Victorian genomes (1.5% vs 0.2%). The Australian genomes also displayed a higher degree of functional similarity, reflected by fewer significantly enriched GO terms relative to each assembly, compared to the Californian assembly. Surprisingly, the Australian giant kelp genomes shared a slightly higher number of orthologous groups with the North American Pacific bull kelp (*N. luetkeana*) than with the Californian giant kelp genome. These findings provide further weight to population genetics studies that have found genetic differences between Northern and Southern hemisphere giant kelp (Assis et al., 2023; Bemmels et al., 2025; Gonzalez et al., 2023), highlighting the need for locally representative genome assemblies (Valiente-Mullor et al., 2021).

The Victorian, Tasmanian and Californian genomes displayed differences in gene ontologies associated with primary metabolism, oxidative stress response and proteostasis, in addition to gene regulation and expression pathways. The enrichment of glycolytic and lipid metabolic processes in the Victorian genome, relative to the Tasmanian genome, suggests functional variation in energy acquisition, storage and membrane composition. Enrichment of proteostasis related processes – protein folding, proteolysis, ubiquitination and chaperone-mediated folding – in the Tasmanian genome suggests enhanced protein regulation mechanisms, essential for maintaining cellular function under conditions that promote protein damage or misfolding (Balchin et al., 2016). The Californian genome was enriched in oxidative stress and redox-related processes, including glutathione metabolism, a pathway central to mitigating reactive oxygen species generated during photosynthesis and environmental stress (Krueger et al., 2014; Noctor et al., 2012). The greater antioxidant capacity may reflect adaptation by the Californian genome to a more variable or stressful environment, compared to the Australian genomes. These differing enrichment patterns may reflect lineage-specific adaptive strategies to grow and survive in their respective environmental conditions. Further functional characterisation using gene transcript evidence is needed to determine whether the observed differences in GO term enrichments translate to functional differences between genomes,.

## Conclusions

We provide two high-quality giant kelp genomes assembled to near chromosome level, which represent the first assemblies from the Southern Hemisphere distribution. We observed a high divergence between Australian and Californian genomes, both at the structural and functional levels. Our findings reinforce the need for multiple, regionally representative genomes for the genomic characterisation of a species. These two high quality genomes from Australian specimens will aid in genomic evolutionary studies of this species and provide the foundation for genome-informed conservation of remnant giant kelp forests around the Southeast Australian shelf.

## Acknowledgments

HS thanks Dr. Carolina Correa-Ospina and Ziyan Zhang from the Biomolecular Resource Facility (Australian National University) for their guidance during the sequencing process. HS also wishes to thank Dr. Rahul Rane and Dr. Gunjan Pandey from the Applied Genomics Initiative (CSIRO) for lending their expertise in genome assembly.

This research was supported by Google Australia.

## Data availability statement

All data needed to evaluate the conclusions in this study are publicly available at CSIRO’s Data Access Portal (https://data.csiro.au/collection/csiro:72991). The raw sequence files are deposited at NCBI’s Sequence Read Archive, under BioProject accession numbers PRJNA1405138 for OTW_2 and PRJNA1357414 for SHL_1. The primary and alternate genome assemblies generated in this study are archived at NCBI’s Genome database, under Genome accession numbers JBTYHO000000000 and JBTYHP000000000 for OTW_2, and JBTYHK000000000 and JBTYHL000000000 for SHL_1.

## References

1. Alexa, A., & Rahnenführer, J. (2025). topGO: Enrichment Analysis for Gene Ontology (R package version 2.58.0). Bioconductor. doi:10.18129/B9.bioc.topGO

2. Alves-Lima, C., Montecinos, G., Escalona, M., Calhoun, S., Marimuthu, M., Nguyen, O., Beraut, E., Lipzen, A., Grigoriev, I. V, Raimondi, P., Nuzhdin, S., Alberto, F., & kelp genome, B. (2025). The reference genome for the northeastern Pacific bull kelp, Nereocystis luetkeana. Journal of Heredity. 10.1093/JHERED/ESAF077

3. Assis, J., Alberto, F., Macaya, E. C., Castilho Coelho, N., Faugeron, S., Pearson, G. A., Ladah, L., Reed, D. C., Raimondi, P., Mansilla, A., Brickle, P., Zuccarello, G. C., & Serrão, E. A. (2023). Past climate-driven range shifts structuring intraspecific biodiversity levels of the giant kelp (Macrocystis pyrifera) at global scales. Scientific Reports 2023 13:1, 13(1), 1–13. 10.1038/s41598-023-38944-7

4. Astashyn, A., Tvedte, E. S., Sweeney, D., Sapojnikov, V., Bouk, N., Joukov, V., Mozes, E., Strope, P. K., Sylla, P. M., Wagner, L., Bidwell, S. L., Brown, L. C., Clark, K., Davis, E. W., Smith-White, B., Hlavina, W., Pruitt, K. D., Schneider, V. A., & Murphy, T. D. (2024). Rapid and sensitive detection of genome contamination at scale with FCS-GX. Genome Biology, 25(1), 1–25. 10.1186/s13059-024-03198-7

5. Balchin, D., Hayer-Hartl, M., & Hartl, F. U. (2016). In vivo aspects of protein folding and quality control. Science, 353(6294). 10.1126/science.aac4354

6. Bemmels, J. B., Starko, S., Weigel, B. L., Hirabayashi, K., Pinch, A., Elphinstone, C., Dethier, M. N., Rieseberg, L. H., Page, J. E., Neufeld, C. J., & Owens, G. L. (2025). Population genomics reveals strong impacts of genetic drift without purging and guides conservation of bull and giant kelp. Current Biology, 35(3), 688–698.e8. 10.1016/j.cub.2024.12.025

7. Brandies, P., Peel, E., Hogg, C. J., & Belov, K. (2019). The Value of Reference Genomes in the Conservation of Threatened Species. Genes 2019, Vol. 10, Page 846, 10(11), 846. 10.3390/genes10110846

8. Brůna, T., Lomsadze, A., & Borodovsky, M. (2024). GeneMark-ETP significantly improves the accuracy of automatic annotation of large eukaryotic genomes. Genome Research, 34(5), 757–768. 10.1101/GR.278373.123/-/DC1

9. Butler, C. L., Lucieer, V. L., Wotherspoon, S. J., & Johnson, C. R. (2020). Multi-decadal decline in cover of giant kelp Macrocystis pyrifera at the southern limit of its Australian range. Marine Ecology Progress Series, 653, 1–18. 10.3354/MEPS13510

10. Chen, J., Zang, Y., Shang, S., & Tang, X. (2019). The complete mitochondrial genome of the brown alga Macrocystis integrifolia (Laminariales, Phaeophyceae). Mitochondrial DNA Part B: Resources, 4(1), 635–636. 10.1080/23802359.2018.1495114

11. Cheng, H., Concepcion, G. T., Feng, X., Zhang, H., & Li, H. (2021). Haplotype-resolved de novo assembly using phased assembly graphs with hifiasm. Nature Methods 2021 18:2, 18(2), 170–175. 10.1038/s41592-020-01056-5

12. Cole, K. (1968). Gametophytic development and fertilization in Macrocystis integrifolia. Canadian Journal of Botany, 46(6), 777–782. 10.1139/B68-108

13. Coombe, L., Kazemi, P., Wong, J., Birol, I., Warren, R. L., & Smith, M. (2024). Multi-genome synteny detection using minimizer graph mappings. BioRxiv, 2024.02.07.579356. 10.1101/2024.02.07.579356

14. Coombe, L., Warren, R. L., & Birol, I. (2025). ntSynt-viz: Visualizing synteny patterns across multiple genomes. BioRxiv, 2025.01.15.633221. 10.1101/2025.01.15.633221

15. Coombe, L., Warren, R. L., Wong, J., Nikolic, V., & Birol, I. (2023). ntLink: A Toolkit for De Novo Genome Assembly Scaffolding and Mapping Using Long Reads. Current Protocols, 3(4), e733. 10.1002/cpz1.733

16. Delehelle, F., Cussat-Blanc, S., Alliot, J. M., Luga, H., & Balaresque, P. (2018). ASGART: fast and parallel genome scale segmental duplications mapping. Bioinformatics, 34(16), 2708–2714. 10.1093/BIOINFORMATICS/BTY172

17. Diesel, J., Molano, G., Montecinos, G. J., DeWeese, K., Calhoun, S., Kuo, A., Lipzen, A., Salamov, A., Grigoriev, I. V., Reed, D. C., Miller, R. J., Nuzhdin, S. V., & Alberto, F. (2023). A scaffolded and annotated reference genome of giant kelp (Macrocystis pyrifera). BMC Genomics, 24(1), 1–13. 10.1186/s12864-023-09658-x

18. Doyle, J. J., & Doyle, J. L. (1987). A rapid DNA isolation procedure for small quantities of fresh leaf tissue. PHYTOCHEMICAL BULLETIN, 19(1), 11. https://worldveg.tind.io/record/33886

19. Dussex, N., van der Valk, T., Morales, H. E., Wheat, C. W., Díez-del-Molino, D., von Seth, J., Foster, Y., Kutschera, V. E., Guschanski, K., Rhie, A., Phillippy, A. M., Korlach, J., Howe, K., Chow, W., Pelan, S., Mendes Damas, J. D., Lewin, H. A., Hastie, A. R., Formenti, G., … Dalén, L. (2021). Population genomics of the critically endangered kākāpō. Cell Genomics, 1(1), 100002. 10.1016/J.XGEN.2021.100002

20. Emms, D. M., Liu, Y., Belcher, L., Holmes, J., & Kelly, S. (2025). OrthoFinder: scalable phylogenetic orthology inference for comparative genomics. BioRxiv, 2025.07.15.664860. 10.1101/2025.07.15.664860

21. Flynn, J. M., Hubley, R., Goubert, C., Rosen, J., Clark, A. G., Feschotte, C., & Smit, A. F. (2020). RepeatModeler2 for automated genomic discovery of transposable element families. Proceedings of the National Academy of Sciences of the United States of America, 117(17), 9451–9457. 10.1073/pnas.1921046117

22. Forbes, H., Visch, W., Bennett, S., Sanderson, J. C., Wright, J. T., & Layton, C. (2025). A historical review of giant kelp harvesting in Tasmania. Journal of Phycology, 61(3), 574–586. 10.1111/jpy.70015

23. Formenti, G., Theissinger, K., Fernandes, C., Bista, I., Bombarely, A., Bleidorn, C., Ciofi, C., Crottini, A., Godoy, J. A., Höglund, J., Malukiewicz, J., Mouton, A., Oomen, R. A., Paez, S., Palsbøll, P. J., Pampoulie, C., Ruiz-López, M. J., Svardal, H., Theofanopoulou, C., … Zammit, G. (2022). The era of reference genomes in conservation genomics. Trends in Ecology & Evolution, 37(3), 197–202. 10.1016/J.TREE.2021.11.008

24. Gabriel, L., Hoff, K. J., Brůna, T., Borodovsky, M., & Stanke, M. (2021). TSEBRA: transcript selector for BRAKER. BMC Bioinformatics, 22(1), 1–12. 10.1186/s12859-021-04482-0

25. Gonzalez, S. T., Alberto, F., & Molano, G. (2023). Whole-genome sequencing distinguishes the two most common giant kelp ecomorphs. Evolution, 77(6), 1354–1369. 10.1093/evolut/qpad093

26. Gonzalez-Aragon, D., Rivadeneira, M. M., Lara, C., Torres, F. I., Vásquez, J. A., & Broitman, B. R. (2024). A species distribution model of the giant kelp Macrocystis pyrifera: Worldwide changes and a focus on the Southeast Pacific. Ecology and Evolution, 14(3), e10901. 10.1002/ECE3.10901

27. Greiner, S., Lehwark, P., & Bock, R. (2019). OrganellarGenomeDRAW (OGDRAW) version 1.3.1: expanded toolkit for the graphical visualization of organellar genomes. Nucleic Acids Research, 47(W1), W59–W64. 10.1093/NAR/GKZ238

28. Guan, D., Guan, D., McCarthy, S. A., Wood, J., Howe, K., Wang, Y., Durbin, R., & Durbin, R. (2020). Identifying and removing haplotypic duplication in primary genome assemblies. Bioinformatics, 36(9), 2896–2898. 10.1093/BIOINFORMATICS/BTAA025

29. Guo, L., Zhong, M., Wang, W., Li, X., & Zhang, Q. (2025). Chromosome-scale genome assembly and annotation of Saccharina sculpera. Scientific Data, 12(1), 1759. 10.1038/S41597-025-06033-1

30. Hoff, K. J., Lange, S., Lomsadze, A., Borodovsky, M., & Stanke, M. (2016). BRAKER1: Unsupervised RNA-Seq-Based Genome Annotation with GeneMark-ET and AUGUSTUS. Bioinformatics, 32(5), 767–769. 10.1093/BIOINFORMATICS/BTV661

31. Hubley, R., Finn, R. D., Clements, J., Eddy, S. R., Jones, T. A., Bao, W., Smit, A. F. A., & Wheeler, T. J. (2016). The Dfam database of repetitive DNA families. Nucleic Acids Research, 44(D1), D81–D89. 10.1093/NAR/GKV1272

32. Iha, C., Layton, C., Flentje, W., Lenton, A., Johnson, C., Fraser, C. I., & Willis, A. (2023). Organellar genomes of giant kelp from the southern hemisphere. Applied Phycology, 4(1), 78–86. 10.1080/26388081.2023.2193619

33. Johnson, C. R., Banks, S. C., Barrett, N. S., Cazassus, F., Dunstan, P. K., Edgar, G. J., Frusher, S. D., Gardner, C., Haddon, M., Helidoniotis, F., Hill, K. L., Holbrook, N. J., Hosie, G. W., Last, P. R., Ling, S. D., Melbourne-Thomas, J., Miller, K., Pecl, G. T., Richardson, A. J., … Taw, N. (2011). Climate change cascades: Shifts in oceanography, species’ ranges and subtidal marine community dynamics in eastern Tasmania. Journal of Experimental Marine Biology and Ecology, 400(1–2), 17–32. 10.1016/J.JEMBE.2011.02.032

34. Johnson, W. E., Koepfl, K., Johnson, W. E., Koepfl, K., & Holt, W. V. (2014). The Role of Genomics in Conservation and Reproductive Sciences. Advances in Experimental Medicine and Biology, 753, 71–96. 10.1007/978-1-4939-0820-2_5

35. Jones, P., Binns, D., Chang, H. Y., Fraser, M., Li, W., McAnulla, C., McWilliam, H., Maslen, J., Mitchell, A., Nuka, G., Pesseat, S., Quinn, A. F., Sangrador-Vegas, A., Scheremetjew, M., Yong, S. Y., Lopez, R., & Hunter, S. (2014). InterProScan 5: genome-scale protein function classification. Bioinformatics, 30(9), 1236–1240. 10.1093/BIOINFORMATICS/BTU031

36. Käll, L., Krogh, A., & Sonnhammer, E. L. L. (2004). A Combined Transmembrane Topology and Signal Peptide Prediction Method. Journal of Molecular Biology, 338(5), 1027–1036. 10.1016/J.JMB.2004.03.016

37. Krogh, A., Larsson, B., Von Heijne, G., & Sonnhammer, E. L. L. (2001). Predicting transmembrane protein topology with a hidden markov model: application to complete genomes. Journal of Molecular Biology, 305(3), 567–580. 10.1006/JMBI.2000.4315

38. Krueger, T., Becker, S., Pontasch, S., Dove, S., Hoegh-Guldberg, O., Leggat, W., Fisher, P. L., & Davy, S. K. (2014). Antioxidant plasticity and thermal sensitivity in four types of Symbiodinium sp. Journal of Phycology, 50(6), 1035–1047. 10.1111/jpy.12232

39. Krzywinski, M., Schein, J., Birol, I., Connors, J., Gascoyne, R., Horsman, D., Jones, S. J., & Marra, M. A. (2009). Circos: An information aesthetic for comparative genomics. Genome Research, 19(9), 1639–1645. 10.1101/GR.092759.109

40. Kuznetsov, D., Tegenfeldt, F., Manni, M., Seppey, M., Berkeley, M., Kriventseva, E. V., & Zdobnov, E. M. (2023). OrthoDB v11: annotation of orthologs in the widest sampling of organismal diversity. Nucleic Acids Research, 51(D1), D445–D451. 10.1093/NAR/GKAC998

41. Laetsch, D. R., & Blaxter, M. L. (2017). BlobTools: Interrogation of genome assemblies. F1000Research 2017 6:1287, 6, 1287. 10.12688/f1000research.12232.1

42. Lewis, R. J., & Neushul, M. (1994). NORTHERN AND SOUTHERN HEMISPHERE HYBRIDS OF MACROCYSTIS (PHAEOPHYCEAE)1. Journal of Phycology, 30(2), 346–353. 10.1111/J.0022-3646.1994.00346.X

43. Li, H. (2018). Minimap2: pairwise alignment for nucleotide sequences. Bioinformatics, 34(18), 3094–3100. 10.1093/BIOINFORMATICS/BTY191

44. Liao, W. W., Asri, M., Ebler, J., Doerr, D., Haukness, M., Hickey, G., Lu, S., Lucas, J. K., Monlong, J., Abel, H. J., Buonaiuto, S., Chang, X. H., Cheng, H., Chu, J., Colonna, V., Eizenga, J. M., Feng, X., Fischer, C., Fulton, R. S., … Paten, B. (2023). A draft human pangenome reference. Nature 2023 617:7960, 617(7960), 312–324. 10.1038/s41586-023-05896-x

45. Ling, S. D. (2008). Range expansion of a habitat-modifying species leads to loss of taxonomic diversity: A new and impoverished reef state. Oecologia, 156(4), 883–894. 10.1007/s00442-008-1043-9

46. Manni, M., Berkeley, M. R., Seppey, M., Simão, F. A., & Zdobnov, E. M. (2021). BUSCO Update: Novel and Streamlined Workflows along with Broader and Deeper Phylogenetic Coverage for Scoring of Eukaryotic, Prokaryotic, and Viral Genomes. Molecular Biology and Evolution, 38(10), 4647–4654. 10.1093/MOLBEV/MSAB199

47. Morin, P. A., Archer, F. I., Avila, C. D., Balacco, J. R., Bukhman, Y. V., Chow, W., Fedrigo, O., Formenti, G., Fronczek, J. A., Fungtammasan, A., Gulland, F. M. D., Haase, B., Peter Heide-Jorgensen, M., Houck, M. L., Howe, K., Misuraca, A. C., Mountcastle, J., Musser, W., Paez, S., … Jarvis, E. D. (2021). Reference genome and demographic history of the most endangered marine mammal, the vaquita. Molecular Ecology Resources, 21(4), 1008–1020. 10.1111/1755-0998.13284;JOURNAL:JOURNAL:14718286;WGROUP:STRING:PUBLICATION

48. Noctor, G., Mhamdi, A., Chaouch, S., Han, Y., Neukermans, J., Marquez-Garcia, B., Queval, G., & Foyer, C. H. (2012). Glutathione in plants: An integrated overview. Plant, Cell and Environment, 35(2), 454–484. 10.1111/J.1365-3040.2011.02400.X;CTYPE:STRING:JOURNAL

49. Oliver, E. C. J., Burrows, M. T., Donat, M. G., Sen Gupta, A., Alexander, L. V., Perkins-Kirkpatrick, S. E., Benthuysen, J. A., Hobday, A. J., Holbrook, N. J., Moore, P. J., Thomsen, M. S., Wernberg, T., & Smale, D. A. (2019). Projected Marine Heatwaves in the 21st Century and the Potential for Ecological Impact. Frontiers in Marine Science, 6, 734. 10.3389/FMARS.2019.00734

50. Oliver, E. C. J., & Holbrook, N. J. (2014). Extending our understanding of South Pacific gyre “spin-up”: Modeling the East Australian Current in a future climate. Journal of Geophysical Research: Oceans, 119(5), 2788–2805. 10.1002/2013JC009591

51. Ondov, B. D., Treangen, T. J., Melsted, P., Mallonee, A. B., Bergman, N. H., Koren, S., & Phillippy, A. M. (2016). Mash: Fast genome and metagenome distance estimation using MinHash. Genome Biology, 17(1), 1–14. 10.1186/S13059-016-0997-X/FIGURES/5

52. Petersen, T. N., Brunak, S., Von Heijne, G., & Nielsen, H. (2011). SignalP 4.0: discriminating signal peptides from transmembrane regions. Nature Methods 2011 8:10, 8(10), 785–786. 10.1038/nmeth.1701

53. Quinlan, A. R., & Hall, I. M. (2010). BEDTools: a flexible suite of utilities for comparing genomic features. Bioinformatics, 26(6), 841–842. 10.1093/BIOINFORMATICS/BTQ033

54. Riehl, K., Riccio, C., Miska, E. A., & Hemberg, M. (2022). TransposonUltimate: software for transposon classification, annotation and detection. Nucleic Acids Research, 50(11), e64. 10.1093/NAR/GKAC136

55. Shan, T., Yuan, J., Su, L., Li, J., Leng, X., Zhang, Y., Gao, H., & Pang, S. (2020). First Genome of the Brown Alga Undaria pinnatifida: Chromosome-Level Assembly Using PacBio and Hi-C Technologies. Frontiers in Genetics, 11. 10.3389/FGENE.2020.00140/FULL

56. Shu, N. (2019). ProtExcluder. GitHub. https://github.com/NBISweden/ProtExcluder

57. Stanke, M., Keller, O., Gunduz, I., Hayes, A., Waack, S., & Morgenstern, B. (2006). AUGUSTUS: ab initio prediction of alternative transcripts. Nucleic Acids Research, 34(suppl_2), W435–W439. 10.1093/NAR/GKL200

58. Starko, S., Bringloe, T. T., Soto Gomez, M., Darby, H., Graham, S. W., & Martone, P. T. (2021). Genomic Rearrangements and Sequence Evolution across Brown Algal Organelles. Genome Biology and Evolution, 13(7). 10.1093/GBE/EVAB124

59. Tarailo-Graovac, M., & Chen, N. (2009). Using RepeatMasker to identify repetitive elements in genomic sequences. Current Protocols in Bioinformatics, 25(25), 4.10.1–4.10.14. 10.1002/0471250953.bi0410s25

60. Thurstan, R. H., Brittain, Z., Jones, D. S., Cameron, E., Dearnaley, J., & Bellgrove, A. (2018). Aboriginal uses of seaweeds in temperate Australia: an archival assessment. Journal of Applied Phycology, 30(3), 1821–1832. 10.1007/S10811-017-1384-Z/FIGURES/1

61. Tillich, M., Lehwark, P., Pellizzer, T., Ulbricht-Jones, E. S., Fischer, A., Bock, R., & Greiner, S. (2017). GeSeq – versatile and accurate annotation of organelle genomes. Nucleic Acids Research, 45(W1), W6–W11. 10.1093/NAR/GKX391

62. Valiente-Mullor, C., Beamud, B., Ansari, I., Frances-Cuesta, C., Garcia-Gonzalez, N., Mejia, L., Ruiz-Hueso, P., & Gonzalez-Candelas, F. (2021). One is not enough: On the effects of reference genome for the mapping and subsequent analyses of short-reads. PLOS Computational Biology, 17(1), e1008678. 10.1371/JOURNAL.PCBI.1008678

63. Walker, B. J., Abeel, T., Shea, T., Priest, M., Abouelliel, A., Sakthikumar, S., Cuomo, C. A., Zeng, Q., Wortman, J., Young, S. K., & Earl, A. M. (2014). Pilon: An Integrated Tool for Comprehensive Microbial Variant Detection and Genome Assembly Improvement. PLOS ONE, 9(11), e112963. 10.1371/JOURNAL.PONE.0112963

64. Walker, F. T. (1952). Chromosome Number of Macrocystis integrifolia Bory. Annals of Botany, 16(1), 23–27. 10.1093/oxfordjournals.aob.a083300

65. Wood, G., Marzinelli, E. M., Vergés, A., Campbell, A. H., Steinberg, P. D., & Coleman, M. A. (2020). Using genomics to design and evaluate the performance of underwater forest restoration. Journal of Applied Ecology, 57(10), 1988–1998. 10.1111/1365-2664.13707

66. Xu, M., Guo, L., Gu, S., Wang, O., Zhang, R., Peters, B. A., Fan, G., Liu, X., Xu, X., Deng, L., & Zhang, Y. (2020). TGS-GapCloser: A fast and accurate gap closer for large genomes with low coverage of error-prone long reads. GigaScience, 9(9), 1–11. 10.1093/GIGASCIENCE/GIAA094

67. Yabu, H., & Sanbonsuga, Y. (1987). Chromosome Count in Macrocystis integrifolia Bory. BULLETIN OF THE FACULTY OF FISHERIES HOKKAIDO UNIVERSITY, 38(4), 339–342.

68. Ye, N., Zhang, X., Miao, M., Fan, X., Zheng, Y., Xu, D., Wang, J., Zhou, L., Wang, D., Gao, Y., Wang, Y., Shi, W., Ji, P., Li, D., Guan, Z., Shao, C., Zhuang, Z., Gao, Z., Qi, J., & Zhao, F. (2015). Saccharina genomes provide novel insight into kelp biology. Nature Communications, 6(1), 1–11. 10.1038/ncomms7986

69. Zhou, C., Brown, M., Blaxter, M., Consortium, T. D. T. of L. P., McCarthy, S. A., & Durbin, R. (2024). Oatk: a de novo assembly tool for complex plant organelle genomes. BioRxiv, 2024.10.23.619857. 10.1101/2024.10.23.619857

