## Supplementary material for "Australian giant kelp genome assemblies show distinct Southern Hemisphere genetics": Fig. S

### Supplementary Information

#### Supplementary Methods

##### DNA extraction protocol

**Buffer preparation**

**1.** Set a water bath or oven to 55°C, used to dissolve the lysis buffer and during lysis.

**2.** Prepare the CTAB Lysis buffer fresh on the day of extraction.

CTAB Lysis buffer recipe (100 ml)

| Reagent | Stock concentration | Amount | Final concentration |
|---|---|---|---|
| Milli-Q water | / | 62 ml | / |
| NaCl | 5 M | 24 ml | 1.2 M |
| EDTA | 0.5 M | 4 ml | 20 mM |
| Tris-HCl | 1 M | 10 ml | 100 mM |
| CTAB | powder | 2 g | 2% (w/v) |
| PVP40 | powder | 2 g | 2% (w/v) |
| PEG 8,000 | powder | 1 g | 1% (w/v) |
| β-mercaptoethanol* | 14.3 M | 1 ml | 1 % (v/v) |
| Proteinase K* | 20 mg/ml | 0.5 ml | 100 µg/ml |

* Add the β-mercaptoethanol (10 µl per ml of lysis buffer) and Proteinase K (5 µl per ml of lysis buffer) prior to starting Step 6.

**3.** Incubate at 55°C and mix the lysis buffer for at least 15 min. Ensure the solution is clear before use. Keep at 55°C until ready to start Step 5.

**Cell lysis**

**4.** Dispense 1.5 ml of CTAB Lysis buffer to a 2 ml Lysis tube. If performing a large prep, then dispense 30 ml of CTAB Lysis buffer to a 50 ml Falcon tube.

**5.** Add the ground giant kelp tissue to the CTAB lysis buffer.

**6.** Incubate the samples at 55°C for 1 hour, shaking at 300 rpm if possible. Invert every 10-15 min.

**7.** Cool tubes for 5 min at RT then centrifuge at 3,000 g for 5 min at room temperature (RT). For large preps, centrifuge at 1,000 g.

**8.** Transfer 1 ml of lysate to a new 2 ml Protein LoBind tube. Avoid carryover of any debris. For large preps, transfer 20 ml of lysate to a new 50 ml Falcon tube.

**<u>Contaminant removal</u>**

**9.** Add an equal volume (1 ml) of Chloroform: Isoamyl (24:1). For large preps, add 20 ml of Chloroform: Isoamyl (24:1).

**10.** Mix by inverting 10-15 times. Let stand at RT for 2-3 min. Centrifuge at 10,000 g for 15 min at 4°C. For large preps, centrifuge at 3,000 g.

**11.** Transfer to a new tube the aqueous phase (top layer), which should be ~900 µl (~15 ml for large preps). Avoid carryover of the organic phase.

**12.** Add 4 µl of RNase A (final concentration: 80 µg/ml). Incubate at 37°C for 30 min. For large preps, add 40 µl of RNase A.

**13.** Re-extract the aqueous phase with an equal volume (900 µl) of Chloroform: Isoamyl (24:1). For large preps, add 15 ml of Chloroform: Isoamyl (24:1).

**14**. Centrifuge at 10,000 g for 15 min at 4°C. For large preps, centrifuge at 3,000 g.

**15.** Transfer 750 µl of supernatant to a 1.5 ml DNA LoBind tube (the supernatant is typically ~900 µl). For large preps, transfer 12.5 ml to a 15 ml Falcon tube.

**16.** Add 0.4 x volume of the supernatant (~300 µl) of 5 M Potassium Acetate. For large preps, add 5 ml of 5 M Potassium Acetate.

**17.** Invert 10 times. Incubate on ice for 20 min.

**18.** Centrifuge at 3,500 g for 45 min at 4°C. Decant ~1 ml of supernatant to a new 1.5 ml DNA LoBind tube. For large preps, decant ~17.5 ml to a 50 ml Falcon tube.

**<u>DNA precipitation and washes</u>**

**19.** Add 0.1 x volume of the supernatant (100 µl) of 3 M Sodium Acetate. For large preps, add 1.75 ml of 3 M Sodium Acetate.

**20.** Add approximately 0.7 x volume of Isopropanol (770 µl). Mix gently, invert 10 times. For large preps, add 13.5 ml of Isopropanol.

**21.** Incubate at RT for 30 min.

**22.** Centrifuge at 10,000 g for 30 min at RT. For large preps, centrifuge at 3,000 g.

**23.** Discard the supernatant. Add 750 µl of 70% Ethanol. Gently mix by inversion. For large preps, add ~10 ml of 70% Ethanol (enough to cover the pellet).

**24.** Centrifuge at 10,000 g for 10 min at RT. For large preps, centrifuge at 3,000 g.

**25.** Remove supernatant.

**26.** Repeat Steps 23-25 once more (i.e., two washes in total).

**27.** Let the pellet dry 3-5 min.

**28.** Resuspend in 50 µl of TE buffer. For large preps, resuspend in 100-200 µl, pending on pellet size.

**29**. Let resuspend overnight.

**<u>DNA cleanup</u>**

The extracted DNA needs to be further cleaned before long-read sequencing as contaminants (primarily polysaccharides) are likely still present after extraction.

**30**. Aliquot 10-30 µg of DNA into a 1.5 mL DNA LoBind tube. Increase the volume to 200 µL with 10 mM Tris-HCl (pH 8).

<u>Note</u>: Volume can exceed 200 µl, maximum of 600 µl due to tube capacity at later steps (ethanol precipitation).

**31.** Add RNAse A (1 µl) and Proteinase K (1 µl) to the solution.

<u>Note</u>: Target concentrations of each enzyme is 100 µg/ml.

**32.** Incubate the samples at 55°C for 20 min, shaking at 400-500 rpm.

**33.** Increase the volume to 600 µl with 10 mM Tris-HCl pH 8 (add 400 µl).

**34.** Add an equal volume of Chloroform: Isoamyl (24:1) (600 µl). Mix by inverting 10-15 times.

**35.** Centrifuge at 10,000 g for 1 min at RT.

**36.** Transfer the upper aqueous phase to a new 1.5 mL DNA LoBind tube.

**37.** Repeat the Chloroform: Isoamyl (24:1) cleanup. Supernatant should be ~600 µl.

**38.** Add 1.5 x volume of 100% Ethanol (~900 µl) and 0.1x volume of 3 M Sodium Acetate (~60 µl).

**39.** Incubate on ice for 1 min.

**40.** Centrifuge at 10,000 g for 1 min at 20°C (or RT).

<u>Note</u>: HMW DNA should pellet easily. If no pellet can be seen, centrifuge for longer. Bringing Ethanol concentration to 2x volume is also an option.

**41.** Carefully decant the supernatant as soon as possible, without disturbing the pellet.

**42.** Add approximately 700 µl of freshly prepared 70% Ethanol.

**43.** Let the pellet soak for 1 min at RT to dissolve excess salts.

**44.** Centrifuge at 10,000 g for 1 min at RT.

**45.** Carefully decant the supernatant as soon as possible, without disturbing the pellet.

**46.** Repeat Steps 42-45 for a second 70% Ethanol wash

**47.** Air-dry the pellet for 5 min.

**48.** Dissolve DNA with 100 µL of 10 mM Tris-HCl (pH 8) or TE buffer.

### Supplementary Figures

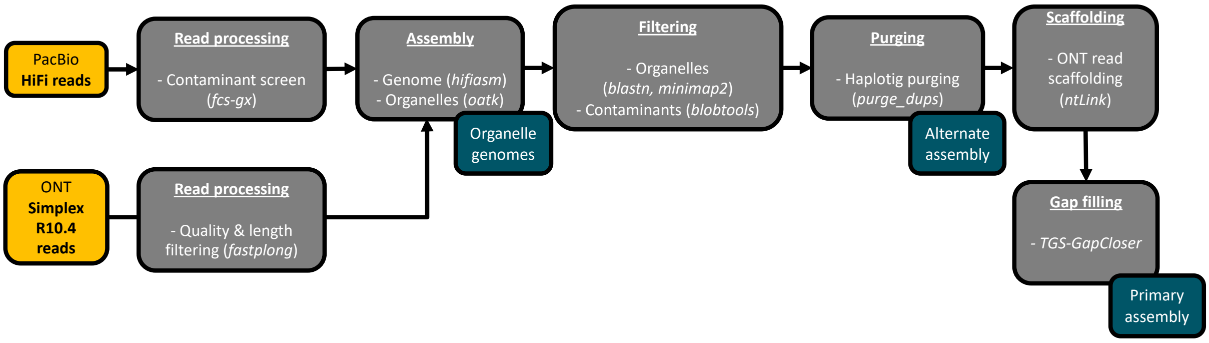


**Figure S1: Overview of the genome assembly pipeline for the Victorian and Tasmanian assemblies.**

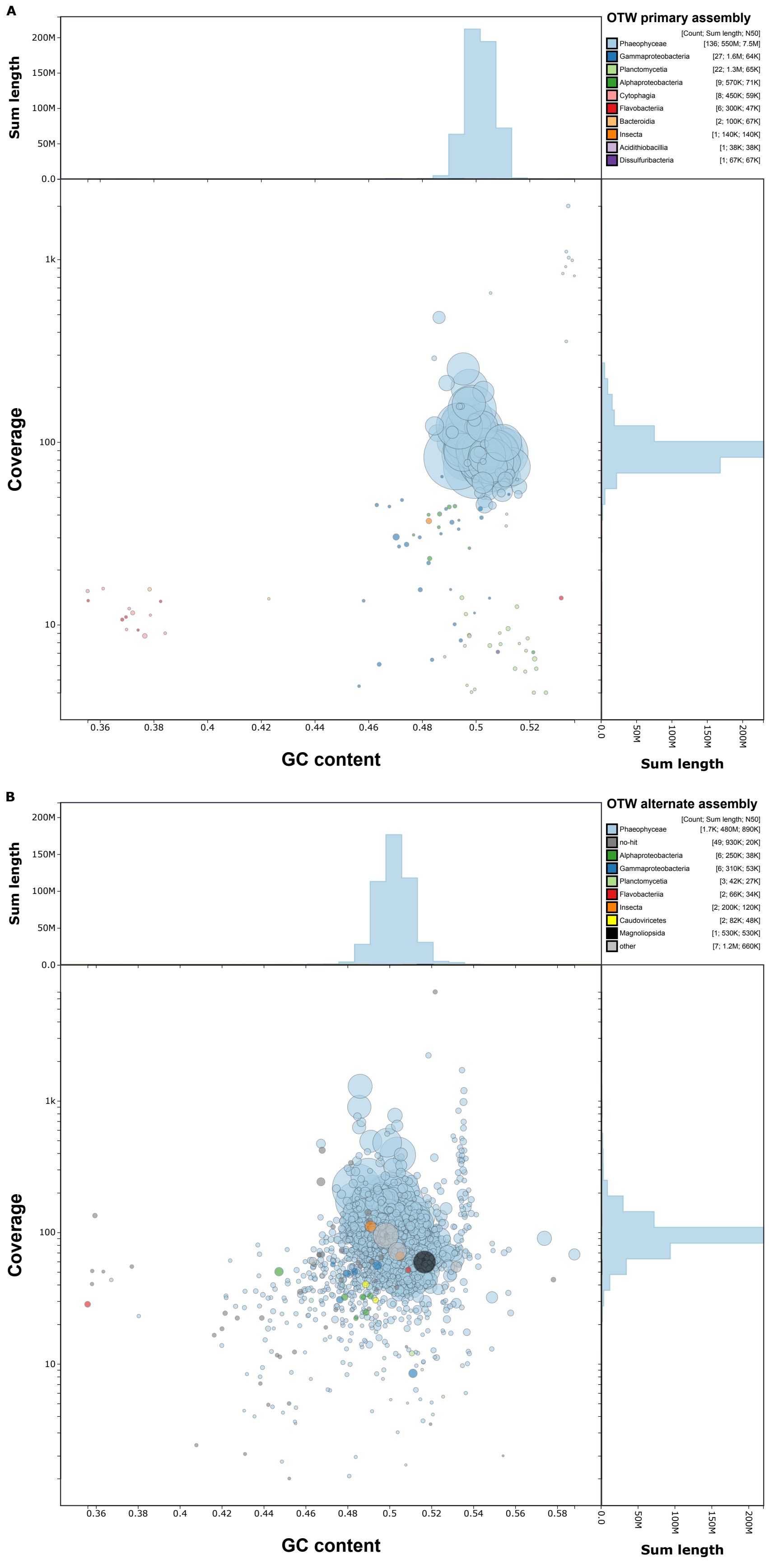


**Figure S2: Blobplots of the primary (A) and alternate (B) genome assemblies for the Victorian sample (OTW).**

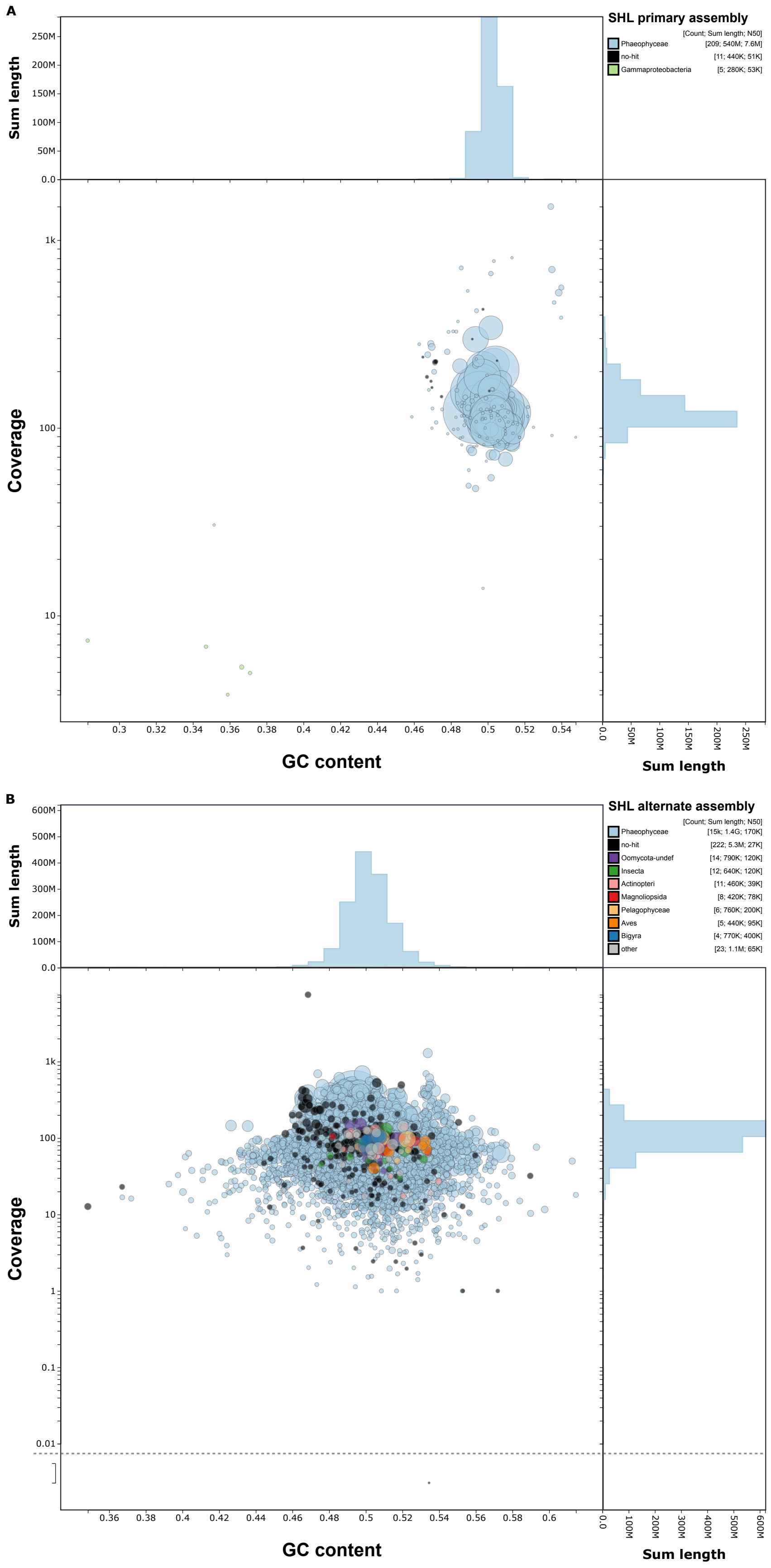


**Figure S3: Blobplots of the primary (A) and alternate (B) genome assemblies for the Tasmanian sample (SHL).**

**A**

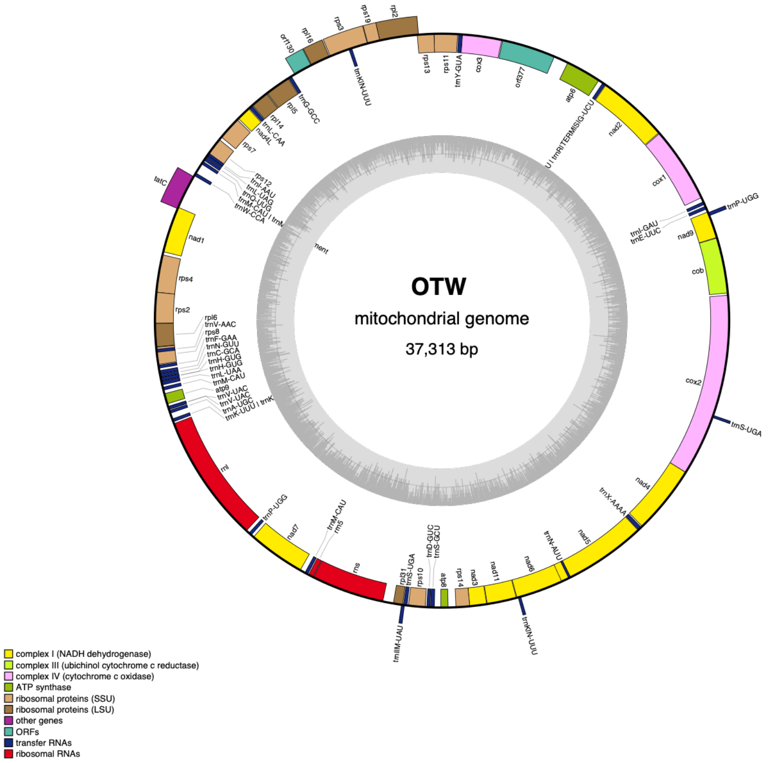


**B**

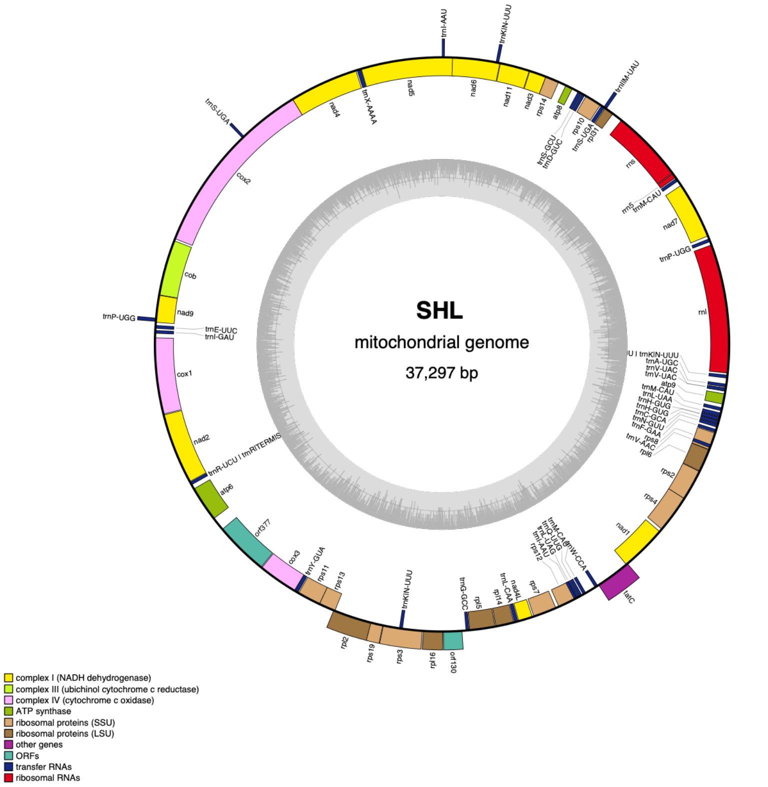


**Figure S4: Mitochondrial genomes from Victorian (A) and Tasmanian (B) samples.**

**A**

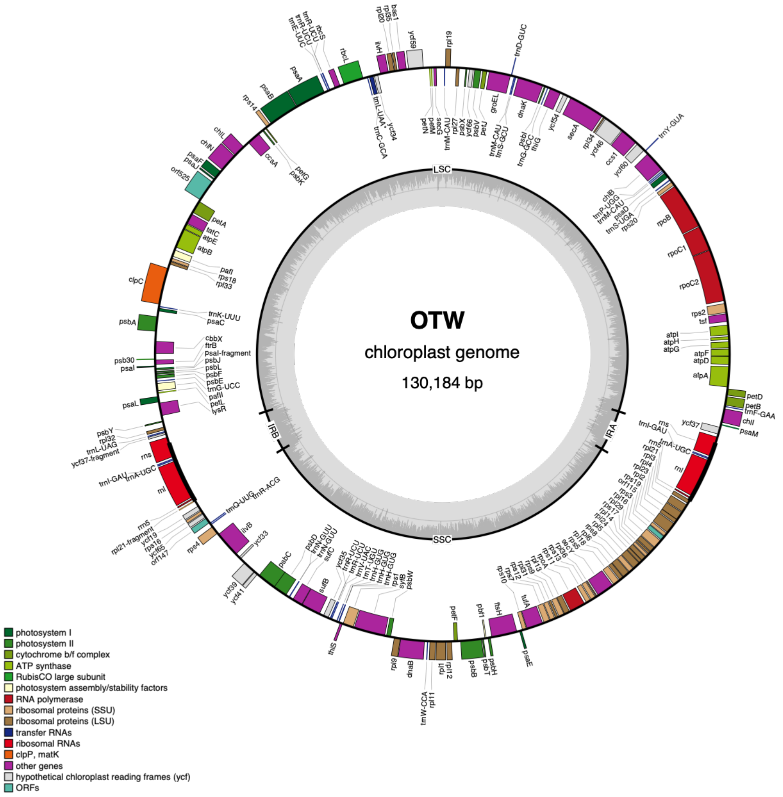


**B**

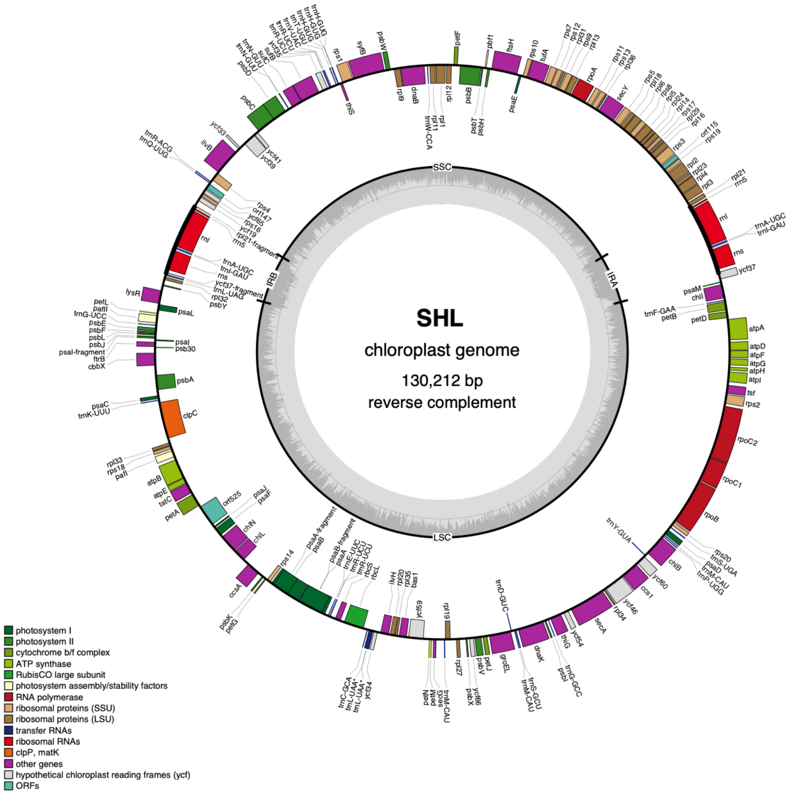


**Figure S5: Chloroplast genomes from Victorian (A) and Tasmanian (B) samples.**

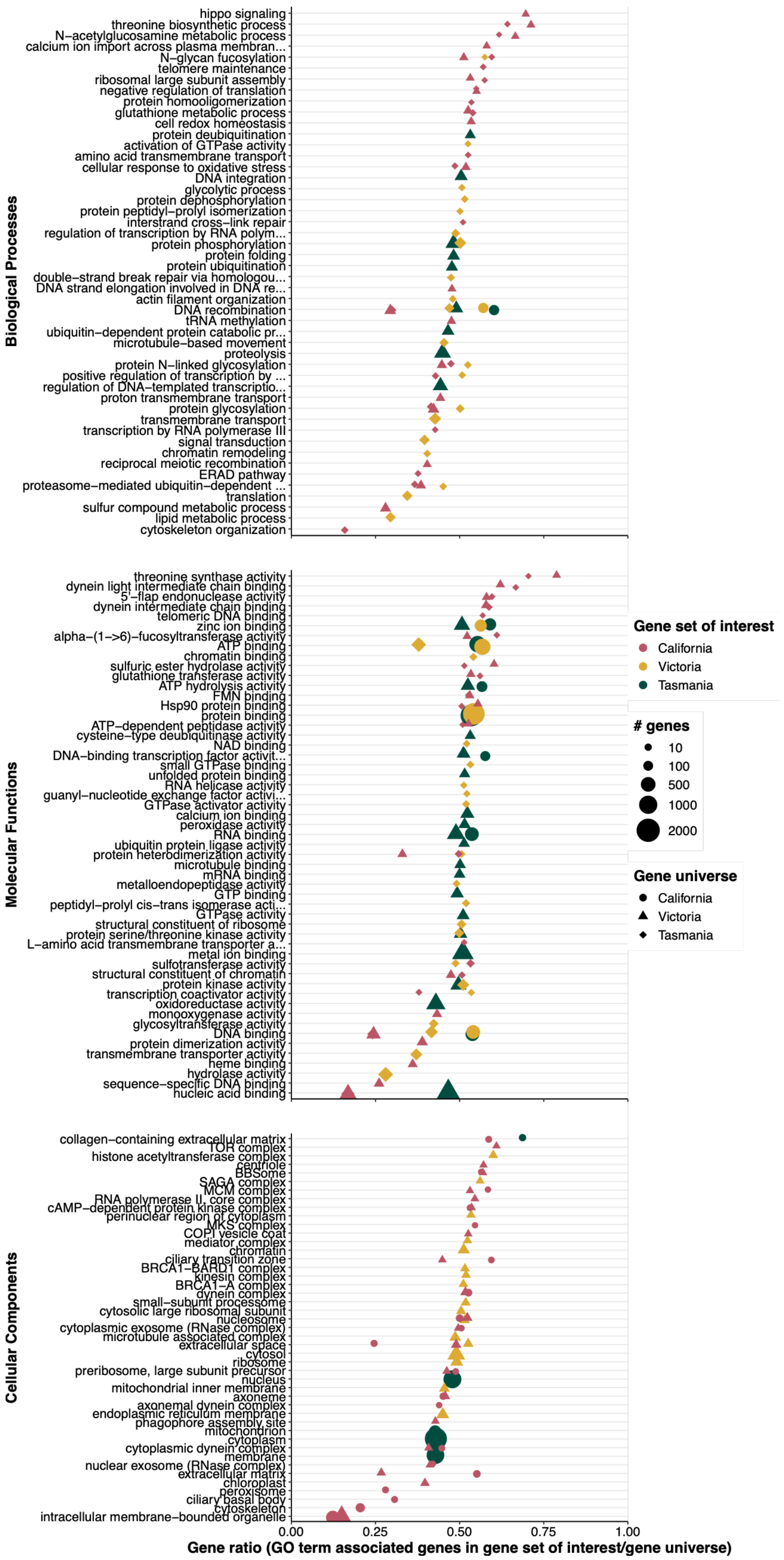


**Figure S6: Enriched gene ontologies (GOs) enriched in Victorian, Tasmanian and Californian giant kelp genomes**. The top 20 significantly enriched GO terms (p<0.01) for each pairwise comparison are shown here.

### Supplementary Tables

**Table S1: Summary of PacBio and ONT sequencing runs for the Victorian (OTW_2) and Tasmanian (SHL_1) samples.**

| Sequence type | Sample ID | Sequencing run | gDNA input (µg) | gDNA median | Size selection threshold | gDNA recovered (%) | Sequencing output summary | | | | Coverage |
|---|---|---|---|---|---|---|---|---|---|---|---|
| | | | | | | | Reads | Yield (Gbp) | Read length, N50 (Kbp) | Read quality, median Q score | |
| PacBio HiFi | OTW_2 | Run 1 | 5.0 | 40.3 | 10 | 16 | 1,273,695 | 15.93 | 14.2 | 33 | 29.7 |
| | OTW_2 | Run 2 | 11.9 | 50.4 | 15 | 33 | 2,264,816 | 28.73 | 14.6 | 34 | 53.5 |
| | OTW_2 | **Combined** | N/A | N/A | N/A | N/A | **3,538,511** | **44.67** | **14.4** | **34** | **83.2** |
| | SHL_1 | Run 1 | 11.9 | 35.5 | 10 | 14 | 2,637,647 | 33.19 | 14.2 | 32 | 61.8 |
| | SHL_1 | Run 2 | 3.5 | 48.7 | 15 | 22 | 2,354,994 | 29.31 | 14.6 | 34 | 54.6 |
| | SHL_1 | **Combined** | N/A | N/A | N/A | N/A | **4,992,641** | **62.50** | **14.4** | **33** | **116.4** |
| ONT R10.4 | OTW_2 | N/A | 5.5 | 24.9 | 15 | 25 | 577,830 | 12.64 | 30.9 | 22 | 23.5 |
| | SHL_1 | N/A | 11.4 | 26.0 | 15 | 28 | 637,730 | 15.30 | 30.9 | 22 | 28.5 |

**Table S2: Summary of sequencing read statistics before (raw reads) and after processing steps.** Statistics of reads used in the genome assembly are indicated in bold. OTW_2 = Victorian sample, SHL_1 = Tasmanian sample.

| Sequence type | Pre-processing step | Sample ID | Reads | Yield (Gbp) | Read length, N50 (Kbp) | Read quality, median Q score | Coverage | Reads removed (%) |
|---|---|---|---|---|---|---|---|---|
| PacBio HiFi | Raw reads | OTW_2 | 3,538,511 | 44.7 | 14.6 | 34 | 83 | N/A |
| | | SHL_1 | 4,992,641 | 62.5 | 14.3 | 33 | 116 | N/A |
| | **Contaminant filtering** | **OTW_2** | **3,473,907** | **43.86** | **14.6** | **34** | **83** | **1.83** |
| | | **SHL_1** | **4,987,661** | **62.44** | **14.3** | **33** | **116** | **0.10** |
| | Raw reads | OTW_2 | 553,642 | 12.15 | 31.1 | 24 | 23 | N/A |
| | | SHL_1 | 603,927 | 14.53 | 31.1 | 24 | 27 | N/A |
| | **Length filtering** | **OTW_2** | **397,173** | **11.46** | **32.4** | 24 | **21** | **28.26** |
| | | **SHL_1** | **491,700** | **14.03** | **31.8** | 24 | **26** | **18.58** |
| **Illumina** | **Raw reads** | **OTW_2** | **162,954,598** | **24.40** | | | **45** | **N/A** |
| | | **SHL_1** | **271,700,226** | **40.60** | | | **76** | **N/A** |

**Table S3: Software and settings used during sequencing read processing, genome assembly and genome annotation.** OTW_2 = Victorian sample, SHL_1 = Tasmanian sample.

| Pipeline | Software | Author | Source | Version | Settings (OTW_2) | Settings (SHL_1) |
|---|---|---|---|---|---|---|
| Read processing | fastplong | Chen *et al.* | https://github.com/OpenGene/fastplong | 0.23.2 | --length_required 10000 | --length_required 10000 |
| | fcs-gx | Astashyn *et al*. | https://github.com/ncbi/fcs?tab=readme-ov-file | 0.5.5 | --mask-transposons=F | --mask-transposons=F |
| Genome assembly | hifiasm | Cheng *et al.* | https://github.com/chhylp123/hifiasm | 0.25.0 | --hg-size 537m<br>-s 0.525<br>--min-hist-cnt 14<br>--hom-cov 92<br>--ul-rate 0.1<br>--ul-cut 10000<br>-k 63<br>-w 63 | --hg-size 537m<br>-s 0.525<br>--min-hist-cnt 9<br>--hom-cov 121<br>--n-hap 3<br>--ul-rate 0.1<br>--ul-cut 10000<br>-k 63<br>-w 63 |
| | oatk | Zhou *et al.* | https://github.com/c-zhou/oatk | 1.0 | default | default |
| | blobtools | Challis *et al.* | https://github.com/genomehubs/blobtoolkit | 4.2.1 | default | default |
| | purge_dups | Guan *et al*. | https://github.com/dfguan/purge_dups | 1.2.5 | -l 5<br>-m 72<br>-h 250<br>-a 95 | -l 5<br>-m 90<br>-h 310<br>-a 95 |
| | ntLink | Coombe *et al.* | https://github.com/bcgsc/ntLink | 1.3.11 | sensitive=True<br>a=2<br>k=32<br>w=500<br>g=10<br>G=10 | sensitive=True<br>a=2<br>k=40<br>w=500<br>g=10<br>G=10 |
| | TGS-GapCloser | Xu *et al.* | https://github.com/BGI-Qingdao/TGS-GapCloser | 1.2.1 | default | default |
| Genome annotation | RepeatMasker | Smit *et al.* | https://github.com/Dfam-consortium/RepeatMasker | 1.4.9 | -s<br>-e rmblast<br>-xsmall | -s<br>-e rmblast<br>-xsmall |
| | BRAKER3 | Gabriel *et al.* | https://github.com/Gaius-Augustus/BRAKER | 3.0.7 | default | default |
| | hisat2 | Kim *et al.* | https://github.com/DaehwanKimLab/hisat2 | 2.2.1 | default | default |
| | GeneMark-ETP | Bruna *et al.* | https://github.com/gatech-genemark/GeneMark-ETP | 1.0.2 | default | default |
| | AUGUSTUS | Stanke *et al.* | https://github.com/Gaius-Augustus/Augustus | 3.4.0 | default | default |
| | TSEBRA | Gabriel et al. | https://github.com/Gaius-Augustus/TSEBRA | 1.1.2.5 | default | default |
| | InterProScan | Jones *et al.* | https://github.com/ebi-pf-team/interproscan | 5.72-103.0 | default | default |

**Table S4: Gene features of mitochondrion assemblies from the Victorian (OTW) and Tasmanian (SHL) samples.**

| Organelle | Gene feature | Gene | Present in |
|---|---|---|---|
| Mitochondrion | tRNA | trnA-UGC | OTW & SHL |
| | | trnR-UCU | OTW & SHL |
| | | trnN-GUU | OTW & SHL |
| | | trnD-GUC | OTW & SHL |
| | | trnC-GCA | OTW & SHL |
| | | trnE-UUC | OTW & SHL |
| | | trnQ-UUG | OTW & SHL |
| | | trnG-GCC | OTW & SHL |
| | | trnH-GUG | OTW & SHL |
| | | trnI-GAU | OTW & SHL |
| | | trnK-UUU | OTW & SHL |
| | | trnL-CAA | OTW & SHL |
| | | trnL-UAG | OTW & SHL |
| | | trnL-UAA | OTW & SHL |
| | | trnM-CAU | OTW & SHL |
| | | trnF-GAA | OTW & SHL |
| | | trnP-UGG | OTW & SHL |
| | | trnS-GCU | OTW & SHL |
| | | trnS-UGA | OTW & SHL |
| | | trnW-CCA | OTW & SHL |
| | | trnY-GUA | OTW & SHL |
| | | trnV-UAC | OTW & SHL |
| | | trnI-AAU | OTW |
| | | trnV-AAC | OTW |
| | | trnN-AUU | OTW |
| | rRNA | rnl | OTW & SHL |
| | | rns | OTW & SHL |
| | | rrn5 | OTW & SHL |
| | ATP synthase | atp6 | OTW & SHL |
| | | atp8 | OTW & SHL |
| | | atp9 | OTW & SHL |
| | Cytochrome oxidase | cox1 | OTW & SHL |
| | | cox2 | OTW & SHL |
| | | cox3 | OTW & SHL |
| | Cytochrome b | cob | OTW & SHL |
| | Maturase | matR | OTW & SHL |
| | NADH dehydrogenase | nad1 | OTW & SHL |
| | | nad2 | OTW & SHL |
| | | nad3 | OTW & SHL |
| | | nad4 | OTW & SHL |
| | | nad5 | OTW & SHL |
| | | nad6 | OTW & SHL |
| | Ribosomal proteins (LSU) | rpl2 | OTW & SHL |
| | | rpl5 | OTW & SHL |
| | | rpl10 | OTW & SHL |
| | | rpl14 | OTW & SHL |
| | | rpl16 | OTW & SHL |
| | | rpl20 | OTW & SHL |
| | | rpl27 | OTW & SHL |
| | | rpl31 | OTW & SHL |
| | Ribosomal proteins (SSU) | rps3 | OTW & SHL |
| | | rps4 | OTW & SHL |
| | | rps7 | OTW & SHL |
| | | rps10 | OTW & SHL |
| | | rps14 | OTW & SHL |
| | | rps12 | OTW |
| | | rps13 | OTW |
| | | rps19 | OTW |
| | Translocon component | tatC | OTW |

**Table S5: Gene features of chloroplast assemblies from the Victorian (OTW) and Tasmanian (SHL) samples.**

| Organelle | Gene feature | Gene | Present in |
|---|---|---|---|
| Chloroplast | tRNA | trnA-UGC | OTW & SHL |
| | | trnC-GCA | OTW & SHL |
| | | trnD-GUC | OTW & SHL |
| | | trnE-UUC | OTW & SHL |
| | | trnF-GAA | OTW & SHL |
| | | trnG-GCC | OTW & SHL |
| | | trnG-UCC | OTW & SHL |
| | | trnH-GUG | OTW & SHL |
| | | trnI-CAU | OTW & SHL |
| | | trnI-GAU | OTW & SHL |
| | | trnK-UUU | OTW & SHL |
| | | trnL-UAG | OTW & SHL |
| | | trnL-UAA | OTW & SHL |
| | | trnM-CAU | OTW & SHL |
| | | trnN-GUU | OTW & SHL |
| | | trnP-UGG | OTW & SHL |
| | | trnQ-UUG | OTW & SHL |
| | | rnR-ACG | OTW & SHL |
| | | trnR-UCU | OTW & SHL |
| | | trnS-GCU | OTW & SHL |
| | | trnS-UGA | OTW & SHL |
| | | trnT-UGU | OTW & SHL |
| | | trnV-GAC | OTW & SHL |
| | | trnV-UAC | OTW & SHL |
| | | trnW-CCA | OTW & SHL |
| | | trnY-GUA | OTW & SHL |
| | | trnL-CAA | SHL |
| | | trnS-GGA | OTW |
| | rRNA | rnl | OTW & SHL |
| | | rns | OTW & SHL |
| | | rrn5 | OTW & SHL |
| | Acetyl-CoA carboxylase | accD | OTW & SHL |
| | ATP synthase | atpA | OTW & SHL |
| | | atpF | OTW & SHL |
| | | atpH | OTW & SHL |
| | | atpI | OTW & SHL |
| | Cellulose synthase | cesA | OTW |
| | Chaperone proteins | dnaK | OTW & SHL |
| | | groEL | OTW & SHL |
| | Cytochrome b6-f complex | petB | OTW & SHL |
| | | petD | OTW & SHL |
| | | petG | OTW & SHL |
| | | petL | OTW & SHL |
| | Ferredoxin-thioredoxin reductase | ftrB | OTW & SHL |
| | Lipopolysaccharide | ftsH | SHL |
| | Maturase | matK | OTW & SHL |
| | Photosystem I | psaE | OTW & SHL |
| | | psaI | OTW & SHL |
| | | psbA | OTW & SHL |
| | | ycf4 | OTW & SHL |
| | | ycf37-fragment | OTW & SHL |
| | | psaD | SHL |
| | | pbf1 | SHL |
| | | ycf3 | SHL |
| | | psaA | OTW |
| | | psaB | OTW |
| | Photosystem II | psbC | OTW & SHL |
| | | psbD | OTW & SHL |
| | | psbE | OTW & SHL |
| | | psbF | OTW & SHL |
| | | psbG | OTW & SHL |
| | | psbJ | OTW & SHL |
| | | psbK | OTW & SHL |
| | | psbL | OTW & SHL |
| | | psbN | OTW & SHL |
| | | psbZ | OTW & SHL |
| | | psb30 | OTW & SHL |
| | | ycf39 | OTW & SHL |
| | Translation initiation factor IF1 | infA | OTW & SHL |
| | Ribosomal Proteins (LSU) | rpl2 | OTW & SHL |
| | | rpl5 | OTW & SHL |
| | | rpl14 | OTW & SHL |
| | | rpl16 | OTW & SHL |
| | | rpl21-fragment | OTW & SHL |
| | | rpl22 | OTW & SHL |
| | | rpl23 | OTW & SHL |
| | | rpl29 | SHL |
| | Ribosomal proteins (SSU) | rps3 | OTW & SHL |
| | | rps7 | OTW & SHL |
| | | rps8 | OTW & SHL |
| | | rps10 | OTW & SHL |
| | | rps11 | OTW & SHL |
| | | rps14 | OTW & SHL |
| | | rps18 | OTW & SHL |
| | | rps19 | OTW & SHL |
| | | rps16 | SHL |
| | | rps20 | SHL |
| | RNA polymerase | rpoB | OTW |
| | | rpoC1 | OTW |
| | | rpoC2 | OTW |
| | RuBisCO | rbcS | OTW |
| | | rbcL | SHL |
| | Other | ycf1 | SHL |
| | | ycf19 | SHL |
| | | ycf34 | SHL |
| | | ycf2 | OTW & SHL |
| | | ycf41 | OTW & SHL |

**Table S6: Summary statistics of the Victorian (OTW_2) and Tasmanian (SHL_1) primary/alternate assemblies after each step of the genome assembly pipeline.**

| Assembly step | Sample | Assembly | Assembly statistics | | | | | |
|---|---|---|---|---|---|---|---|---|
| | | | Size (bp) | Contigs/scaffolds | Largest contig | N50 | N90 | Gaps |
| Assembly | OTW_2 | Primary | **556,791,888** | 311 | 17,810,277 | **7,072,331** | **2,261,158** | - |
| | | Alternate | **484,826,454** | 1,738 | 4,412,481 | **874,697** | **147,894** | - |
| | SHL_1 | Primary | **557,493,934** | 549 | 24,206,539 | **7,417,796** | **2,423,868** | - |
| | | Alternate | **1,435,104,902** | 15,781 | 4,389,366 | **172,024** | **35,436** | - |
| Filtering (organelles) | OTW_2 | Primary | **551,691,849** | 213 | 17,810,277 | **7,072,331** | **2,328,423** | - |
| | | Alternate | **484,625,617** | 1,733 | 4,412,481 | **874,697** | **150,136** | - |
| | SHL_1 | Primary | **538,194,157** | 225 | 24,206,539 | **7,558,681** | **2,951,805** | - |
| | | Alternate | **1,432,207,370** | 15,698 | 4,389,366 | **172,618** | **35,533** | - |
| Filtering (contaminants) | OTW_2 | Primary | **547,048,901** | 136 | 17,810,277 | **7,459,571** | **2,624,379** | - |
| | | Alternate | **482,189,867** | 1,699 | 4,412,481 | **882,897** | **156,653** | - |
| | SHL_1 | Primary | **537,889,544** | 219 | 24,206,539 | **7,558,681** | **2,951,805** | - |
| | | Alternate | **1,427,499,246** | 15,611 | 4,389,366 | **173,277** | **35,620** | - |
| Purging | OTW_2 | Primary | **534,376,919** | 117 | 17,810,277 | **7,702,983** | **2,957,003** | - |
| | | Alternate | **473,362,647** | 1,188 | 4,412,481 | **929,584** | **207,160** | - |
| | SHL_1 | Primary | **527,935,079** | 106 | 24,206,539 | **7,653,012** | **3,155,944** | - |
| | | Alternate | **523,139,414** | 3,241 | 4,389,366 | **332,743** | **73,644** | - |
| Scaffolding | OTW_2 | Primary | **534,372,642** | 39 | 25,450,493 | **15,239,920** | **11,904,287** | 76 |
| | SHL_1 | Primary | **527,935,520** | 49 | 27,845,607 | **15,032,638** | **10,975,686** | 56 |
| Gap filling | OTW_2 | Primary | **534,574,384** | 43 | 25,451,472 | **14,545,337** | **10,622,564** | 56 |
| | SHL_1 | Primary | **527,958,888** | 49 | 27,845,607 | **15,032,638** | **10,975,686** | 47 |

**Table S7: Summary of the repetitive elements identified by *RepeatMasker* and *RepeatModeller* in the Victorian (OTW_2) and Tasmanian (SHL_1) genomes.**

| Type of repetitive element | | | Number of repetitive elements | | Length of repetitive elements (bp) | | Genome proportion (%) | |
|---|---|---|---|---|---|---|---|---|
| | | | OTW_2 | SHL_1 | OTW_2 | SHL_1 | OTW_2 | SHL_1 |
| **Retroelements** | | | **376,955** | **454,948** | **111,746,597** | **124,185,812** | **20.90** | **23.52** |
| | SINEs | | 0 | 1,199 | 0 | 232,920 | 0.00 | 0.04 |
| | Penelope | | 0 | 0 | 0 | 0 | 0.00 | 0.00 |
| | LINEs | | 179,277 | 126,585 | 43,951,213 | 42,972,603 | 8.22 | 8.14 |
| | | CRE/SLACS | 36 | 2,740 | 1,294 | 1,907,516 | 0.00 | 0.36 |
| | | L2/CR1/Rex | 0 | 22,615 | 0 | 4,191,281 | 0.00 | 0.79 |
| | | R1/LOA/Jockey | 56 | 0 | 7,093 | 0 | 0.00 | 0.00 |
| | | R2/R4/NeSL | 0 | 0 | 0 | 0 | 0.00 | 0.00 |
| | | RTE/Bov-B | 159,640 | 101,204 | 41,582,662 | 36,863,336 | 7.78 | 6.98 |
| | | L1/CIN4 | 41 | 0 | 1,472 | 0 | 0.00 | 0.00 |
| | LTR elements | | 197,678 | 327,164 | 67,795,384 | 80,980,289 | 12.68 | 15.34 |
| | | BEL/Pao | 1,442 | 4,756 | 178,200 | 1,538,327 | 0.03 | 0.29 |
| | | Ty1/Copia | 31,520 | 46,171 | 14,892,928 | 13,834,599 | 2.79 | 2.62 |
| | | Gypsy/DIRS1 | 56,352 | 54,184 | 28,345,172 | 29,012,582 | 5.30 | 5.50 |
| | | Retroviral | 0 | 594 | 0 | 136,893 | 0.00 | 0.03 |
| **DNA transposons** | | | **13,401** | **7,651** | **3,480,746** | **2,794,050** | **0.65** | **0.53** |
| | hobo-Activator | | 2,695 | 1,425 | 456,767 | 212,770 | 0.09 | 0.04 |
| | Tc1-IS630-Pogo | | 306 | 511 | 140,849 | 144,734 | 0.03 | 0.03 |
| | En-Spm | | 0 | 0 | 0 | 0 | 0.00 | 0.00 |
| | MULE-MuDR | | 564 | 228 | 153,564 | 126,987 | 0.03 | 0.02 |
| | PiggyBac | | 0 | 0 | 0 | 0 | 0.00 | 0.00 |
| | Tourist/Harbinger | | 4,993 | 884 | 839,005 | 237,498 | 0.16 | 0.04 |
| | Other (Mirage, P-element,Transib) | | 0 | 0 | 0 | 0 | 0.00 | 0.00 |
| **Rolling-circles** | | | 8,097 | 519 | 4,067,937 | 382,216 | **0.76** | **0.07** |
| **Unclassified** | | | 793,068 | 758,973 | 140,496,017 | 126,351,773 | **26.28** | **23.93** |
| **Total interspersed repeats** | | | | | 255,723,360 | 253,331,635 | **47.84** | **47.98** |
| **Small RNA** | | | 4,194 | 3,454 | 757,043 | 855,672 | **0.14** | **0.16** |
| **Satellites** | | | 114 | 0 | 10,277 | 0 | **0.00** | **0.00** |
| **Simple repeats** | | | 597,384 | 595,310 | 34,772,735 | 34,688,154 | **6.50** | **6.57** |
| **Low complexity** | | | 70,924 | 71,050 | 4,526,169 | 4,551,229 | **0.85** | **0.86** |
| **TOTAL** | | | | | **299,857,521** | **293,808,906** | **56.09** | **55.65** |

**Table S8: Significantly enriched gene ontologies (GOs) for the Victorian (OTW), Tasmanian (SHL) and Californian (CAL) giant kelp assemblies.** The pairwise comparison indicates the gene set of interest (e.g., SHL) and the gene universe (e.g., CAL). The gene universe also includes the gene set of interest (i.e., SHL + CAL in this example).

| Ontology | GO ID | Term | Genes - universe | Genes - gene set of interest | Expected | p-value | Gene set of interest | Gene universe | Pairwise comparison |
|---|---|---|---|---|---|---|---|---|---|
| BP | GO:0006310 | DNA recombination | 288 | 170 | 147.51 | 0.0015 | SHL | CAL | SHL/CAL |
| BP | GO:0006468 | protein phosphorylation | 642 | 311 | 255.21 | 3.50E-07 | SHL | OTW | SHL/OTW |
| BP | GO:0006355 | regulation of DNA-templated transcriptio... | 574 | 258 | 228.18 | 6.50E-06 | SHL | OTW | SHL/OTW |
| BP | GO:0006508 | proteolysis | 788 | 364 | 313.25 | 5.20E-05 | SHL | OTW | SHL/OTW |
| BP | GO:0015074 | DNA integration | 271 | 138 | 107.73 | 0.00011 | SHL | OTW | SHL/OTW |
| BP | GO:0006457 | protein folding | 251 | 123 | 99.78 | 0.00182 | SHL | OTW | SHL/OTW |
| BP | GO:0016579 | protein deubiquitination | 118 | 62 | 46.91 | 0.00315 | SHL | OTW | SHL/OTW |
| BP | GO:0016567 | protein ubiquitination | 208 | 101 | 82.68 | 0.00406 | SHL | OTW | SHL/OTW |
| BP | GO:0006310 | DNA recombination | 340 | 169 | 135.16 | 0.00741 | SHL | OTW | SHL/OTW |
| BP | GO:0006511 | ubiquitin-dependent protein catabolic pr... | 265 | 123 | 105.34 | 0.00922 | SHL | OTW | SHL/OTW |
| BP | GO:0006310 | DNA recombination | 286 | 167 | 146.17 | 0.00064 | OTW | CAL | OTW/CAL |
| BP | GO:0006468 | protein phosphorylation | 642 | 321 | 207.63 | 1.70E-20 | OTW | SHL | OTW/SHL |
| BP | GO:0055085 | transmembrane transport | 782 | 333 | 252.9 | 1.20E-13 | OTW | SHL | OTW/SHL |
| BP | GO:0006412 | translation | 536 | 188 | 173.35 | 4.60E-08 | OTW | SHL | OTW/SHL |
| BP | GO:0006310 | DNA recombination | 340 | 156 | 109.96 | 3.50E-07 | OTW | SHL | OTW/SHL |
| BP | GO:0007018 | microtubule-based movement | 242 | 112 | 78.26 | 1.50E-06 | OTW | SHL | OTW/SHL |
| BP | GO:0006357 | regulation of transcription by RNA polym... | 200 | 99 | 64.68 | 0.00013 | OTW | SHL | OTW/SHL |
| BP | GO:0006470 | protein dephosphorylation | 93 | 47 | 30.08 | 2.00E-04 | OTW | SHL | OTW/SHL |
| BP | GO:0000724 | double-strand break repair via homologou... | 112 | 54 | 36.22 | 0.00023 | OTW | SHL | OTW/SHL |
| BP | GO:0006487 | protein N-linked glycosylation | 88 | 45 | 28.46 | 0.00026 | OTW | SHL | OTW/SHL |
| BP | GO:0006629 | lipid metabolic process | 670 | 196 | 216.68 | 0.00031 | OTW | SHL | OTW/SHL |
| BP | GO:0007165 | signal transduction | 475 | 192 | 153.62 | 0.00036 | OTW | SHL | OTW/SHL |
| BP | GO:0045944 | positive regulation of transcription by ... | 87 | 43 | 28.14 | 0.00072 | OTW | SHL | OTW/SHL |
| BP | GO:0006096 | glycolytic process | 69 | 35 | 22.32 | 0.00113 | OTW | SHL | OTW/SHL |
| BP | GO:0000413 | protein peptidyl-prolyl isomerization | 68 | 34 | 21.99 | 0.00182 | OTW | SHL | OTW/SHL |
| BP | GO:0036071 | N-glycan fucosylation | 31 | 18 | 10.03 | 0.00275 | OTW | SHL | OTW/SHL |
| BP | GO:0043161 | proteasome-mediated ubiquitin-dependent ... | 103 | 45 | 33.31 | 0.00283 | OTW | SHL | OTW/SHL |
| BP | GO:0006486 | protein glycosylation | 181 | 90 | 58.54 | 0.00297 | OTW | SHL | OTW/SHL |
| BP | GO:0007015 | actin filament organization | 98 | 47 | 31.69 | 0.00312 | OTW | SHL | OTW/SHL |
| BP | GO:0006338 | chromatin remodeling | 119 | 47 | 38.49 | 0.00374 | OTW | SHL | OTW/SHL |
| BP | GO:0090630 | activation of GTPase activity | 46 | 24 | 14.88 | 0.0041 | OTW | SHL | OTW/SHL |
| BP | GO:0006487 | protein N-linked glycosylation | 89 | 41 | 12.72 | 4.50E-15 | CAL | SHL | CAL/SHL |
| BP | GO:0006749 | glutathione metabolic process | 62 | 33 | 8.86 | 5.70E-13 | CAL | SHL | CAL/SHL |
| BP | GO:0006486 | protein glycosylation | 178 | 76 | 25.44 | 8.30E-12 | CAL | SHL | CAL/SHL |
| BP | GO:0036071 | N-glycan fucosylation | 33 | 20 | 4.72 | 9.90E-10 | CAL | SHL | CAL/SHL |
| BP | GO:0034599 | cellular response to oxidative stress | 52 | 26 | 7.43 | 1.00E-09 | CAL | SHL | CAL/SHL |
| BP | GO:0045944 | positive regulation of transcription by ... | 82 | 34 | 11.72 | 1.60E-09 | CAL | SHL | CAL/SHL |
| BP | GO:0043161 | proteasome-mediated ubiquitin-dependent ... | 103 | 39 | 14.72 | 3.30E-09 | CAL | SHL | CAL/SHL |
| BP | GO:0006310 | DNA recombination | 288 | 86 | 41.16 | 1.20E-08 | CAL | SHL | CAL/SHL |
| BP | GO:0009088 | threonine biosynthetic process | 23 | 15 | 3.29 | 3.10E-08 | CAL | SHL | CAL/SHL |
| BP | GO:0003333 | amino acid transmembrane transport | 33 | 17 | 4.72 | 4.70E-07 | CAL | SHL | CAL/SHL |
| BP | GO:0000027 | ribosomal large subunit assembly | 23 | 13 | 3.29 | 2.80E-06 | CAL | SHL | CAL/SHL |
| BP | GO:0036503 | ERAD pathway | 60 | 23 | 8.58 | 3.50E-06 | CAL | SHL | CAL/SHL |
| BP | GO:0000723 | telomere maintenance | 32 | 18 | 4.57 | 5.60E-06 | CAL | SHL | CAL/SHL |
| BP | GO:0051260 | protein homooligomerization | 22 | 12 | 3.14 | 1.10E-05 | CAL | SHL | CAL/SHL |
| BP | GO:0006044 | N-acetylglucosamine metabolic process | 16 | 10 | 2.29 | 1.20E-05 | CAL | SHL | CAL/SHL |
| BP | GO:0036297 | interstrand cross-link repair | 26 | 13 | 3.72 | 1.70E-05 | CAL | SHL | CAL/SHL |
| BP | GO:0017148 | negative regulation of translation | 20 | 11 | 2.86 | 2.40E-05 | CAL | SHL | CAL/SHL |
| BP | GO:0007010 | cytoskeleton organization | 257 | 42 | 36.73 | 3.30E-05 | CAL | SHL | CAL/SHL |
| BP | GO:0006383 | transcription by RNA polymerase III | 46 | 19 | 6.57 | 3.40E-05 | CAL | SHL | CAL/SHL |
| BP | GO:0006487 | protein N-linked glycosylation | 95 | 42 | 12.92 | 1.80E-14 | CAL | OTW | CAL/OTW |
| BP | GO:0006749 | glutathione metabolic process | 62 | 33 | 8.43 | 1.30E-13 | CAL | OTW | CAL/OTW |
| BP | GO:0006486 | protein glycosylation | 185 | 77 | 25.16 | 2.10E-12 | CAL | OTW | CAL/OTW |
| BP | GO:0034599 | cellular response to oxidative stress | 50 | 26 | 6.8 | 1.10E-10 | CAL | OTW | CAL/OTW |
| BP | GO:0043161 | proteasome-mediated ubiquitin-dependent ... | 102 | 39 | 13.87 | 1.30E-10 | CAL | OTW | CAL/OTW |
| BP | GO:1902600 | proton transmembrane transport | 68 | 30 | 9.25 | 7.20E-10 | CAL | OTW | CAL/OTW |
| BP | GO:0009088 | threonine biosynthetic process | 21 | 15 | 2.86 | 2.30E-09 | CAL | OTW | CAL/OTW |
| BP | GO:0006310 | DNA recombination | 286 | 87 | 38.89 | 5.40E-09 | CAL | OTW | CAL/OTW |
| BP | GO:0030488 | tRNA methylation | 51 | 24 | 6.94 | 7.60E-09 | CAL | OTW | CAL/OTW |
| BP | GO:0036071 | N-glycan fucosylation | 38 | 20 | 5.17 | 1.20E-08 | CAL | OTW | CAL/OTW |
| BP | GO:0045454 | cell redox homeostasis | 36 | 19 | 4.9 | 2.60E-08 | CAL | OTW | CAL/OTW |
| BP | GO:0006790 | sulfur compound metabolic process | 214 | 63 | 29.1 | 5.70E-07 | CAL | OTW | CAL/OTW |
| BP | GO:0000027 | ribosomal large subunit assembly | 24 | 13 | 3.26 | 3.00E-06 | CAL | OTW | CAL/OTW |
| BP | GO:0006044 | N-acetylglucosamine metabolic process | 15 | 10 | 2.04 | 3.30E-06 | CAL | OTW | CAL/OTW |
| BP | GO:0007131 | reciprocal meiotic recombination | 41 | 16 | 5.58 | 5.40E-06 | CAL | OTW | CAL/OTW |
| BP | GO:0035329 | hippo signaling | 13 | 9 | 1.77 | 6.60E-06 | CAL | OTW | CAL/OTW |
| BP | GO:0017148 | negative regulation of translation | 20 | 11 | 2.72 | 1.50E-05 | CAL | OTW | CAL/OTW |
| BP | GO:0006271 | DNA strand elongation involved in DNA re... | 27 | 13 | 3.67 | 1.60E-05 | CAL | OTW | CAL/OTW |
| BP | GO:0098703 | calcium ion import across plasma membran... | 17 | 10 | 2.31 | 1.60E-05 | CAL | OTW | CAL/OTW |
| CC | GO:0005634 | nucleus | 2314 | 1077 | 975.75 | 3.60E-12 | SHL | OTW | SHL/OTW |
| CC | GO:0005737 | cytoplasm | 4446 | 1902 | 1874.76 | 1.70E-08 | SHL | OTW | SHL/OTW |
| CC | GO:0016020 | membrane | 2477 | 1039 | 1044.49 | 1.90E-07 | SHL | OTW | SHL/OTW |
| CC | GO:0005739 | mitochondrion | 746 | 324 | 314.57 | 0.00019 | SHL | OTW | SHL/OTW |
| CC | GO:0062023 | collagen-containing extracellular matrix | 40 | 27 | 16.87 | 0.00105 | SHL | OTW | SHL/OTW |
| CC | GO:0005829 | cytosol | 872 | 428 | 271.81 | 1.00E-27 | OTW | SHL | OTW/SHL |
| CC | GO:0005789 | endoplasmic reticulum membrane | 286 | 127 | 89.15 | 5.70E-07 | OTW | SHL | OTW/SHL |
| CC | GO:0005615 | extracellular space | 94 | 49 | 29.3 | 1.70E-05 | OTW | SHL | OTW/SHL |
| CC | GO:0005840 | ribosome | 267 | 131 | 83.23 | 3.70E-05 | OTW | SHL | OTW/SHL |
| CC | GO:0005743 | mitochondrial inner membrane | 121 | 54 | 37.72 | 0.00014 | OTW | SHL | OTW/SHL |
| CC | GO:0000786 | nucleosome | 75 | 38 | 23.38 | 0.00032 | OTW | SHL | OTW/SHL |
| CC | GO:0022625 | cytosolic large ribosomal subunit | 69 | 35 | 21.51 | 0.00053 | OTW | SHL | OTW/SHL |
| CC | GO:0032040 | small-subunit processome | 67 | 34 | 20.88 | 0.00063 | OTW | SHL | OTW/SHL |
| CC | GO:0000785 | chromatin | 189 | 98 | 58.91 | 0.00145 | OTW | SHL | OTW/SHL |
| CC | GO:0048471 | perinuclear region of cytoplasm | 39 | 21 | 12.16 | 0.00265 | OTW | SHL | OTW/SHL |
| CC | GO:0070531 | BRCA1-A complex | 49 | 25 | 15.27 | 0.00289 | OTW | SHL | OTW/SHL |
| CC | GO:0005871 | kinesin complex | 49 | 25 | 15.27 | 0.00289 | OTW | SHL | OTW/SHL |
| CC | GO:0031436 | BRCA1-BARD1 complex | 45 | 23 | 14.03 | 0.00408 | OTW | SHL | OTW/SHL |
| CC | GO:0016592 | mediator complex | 38 | 20 | 11.85 | 0.00471 | OTW | SHL | OTW/SHL |
| CC | GO:0000124 | SAGA complex | 25 | 14 | 7.79 | 0.00862 | OTW | SHL | OTW/SHL |
| CC | GO:0005875 | microtubule associated complex | 157 | 78 | 48.94 | 0.00891 | OTW | SHL | OTW/SHL |
| CC | GO:0000123 | histone acetyltransferase complex | 51 | 30 | 15.9 | 0.00929 | OTW | SHL | OTW/SHL |

OFFICIAL

| | | | | | | | | | |
|---|---|---|---|---|---|---|---|---|---|
| CC | GO:0005615 | extracellular space | 91 | 45 | 12.19 | 9.60E-17 | CAL | SHL | CAL/SHL |
| CC | GO:0000786 | nucleosome | 76 | 39 | 10.18 | 2.20E-15 | CAL | SHL | CAL/SHL |
| CC | GO:0030286 | dynein complex | 81 | 41 | 10.85 | 7.20E-11 | CAL | SHL | CAL/SHL |
| CC | GO:0043231 | intracellular membrane-bounded organelle | 3812 | 513 | 510.68 | 3.70E-10 | CAL | SHL | CAL/SHL |
| CC | GO:0031012 | extracellular matrix | 83 | 22 | 11.12 | 1.30E-09 | CAL | SHL | CAL/SHL |
| CC | GO:0005930 | axoneme | 67 | 30 | 8.98 | 6.30E-08 | CAL | SHL | CAL/SHL |
| CC | GO:0009507 | chloroplast | 62 | 25 | 8.31 | 1.20E-07 | CAL | SHL | CAL/SHL |
| CC | GO:0005952 | cAMP-dependent protein kinase complex | 24 | 13 | 3.22 | 2.50E-06 | CAL | SHL | CAL/SHL |
| CC | GO:0030687 | preribosome, large subunit precursor | 30 | 14 | 4.02 | 9.90E-06 | CAL | SHL | CAL/SHL |
| CC | GO:0005665 | RNA polymerase II, core complex | 20 | 11 | 2.68 | 1.30E-05 | CAL | SHL | CAL/SHL |
| CC | GO:0005868 | cytoplasmic dynein complex | 37 | 15 | 4.96 | 3.80E-05 | CAL | SHL | CAL/SHL |
| CC | GO:0000177 | cytoplasmic exosome (RNase complex) | 20 | 10 | 2.68 | 9.20E-05 | CAL | SHL | CAL/SHL |
| CC | GO:0030126 | COPI vesicle coat | 17 | 9 | 2.28 | 0.00012 | CAL | SHL | CAL/SHL |
| CC | GO:0005814 | centriole | 14 | 8 | 1.88 | 0.00014 | CAL | SHL | CAL/SHL |
| CC | GO:0034464 | BBSome | 14 | 8 | 1.88 | 0.00014 | CAL | SHL | CAL/SHL |
| CC | GO:0035869 | ciliary transition zone | 25 | 11 | 3.35 | 0.00015 | CAL | SHL | CAL/SHL |
| CC | GO:0000407 | phagophore assembly site | 25 | 11 | 3.35 | 0.00017 | CAL | SHL | CAL/SHL |
| CC | GO:0042555 | MCM complex | 15 | 8 | 2.01 | 0.00027 | CAL | SHL | CAL/SHL |
| CC | GO:0000176 | nuclear exosome (RNase complex) | 24 | 10 | 3.22 | 0.00059 | CAL | SHL | CAL/SHL |
| CC | GO:0038201 | TOR complex | 10 | 6 | 1.34 | 0.00074 | CAL | SHL | CAL/SHL |
| CC | GO:0000786 | nucleosome | 77 | 39 | 9.75 | 5.80E-16 | CAL | OTW | CAL/OTW |
| CC | GO:0043231 | intracellular membrane-bounded organelle | 3773 | 468 | 477.9 | 1.70E-10 | CAL | OTW | CAL/OTW |
| CC | GO:0030286 | dynein complex | 82 | 42 | 10.39 | 1.70E-10 | CAL | OTW | CAL/OTW |
| CC | GO:0031012 | extracellular matrix | 71 | 39 | 8.99 | 1.10E-09 | CAL | OTW | CAL/OTW |
| CC | GO:0062023 | collagen-containing extracellular matrix | 31 | 18 | 3.93 | 2.50E-09 | CAL | OTW | CAL/OTW |
| CC | GO:0005930 | axoneme | 68 | 30 | 8.61 | 3.50E-08 | CAL | OTW | CAL/OTW |
| CC | GO:0005777 | peroxisome | 65 | 19 | 8.23 | 9.40E-07 | CAL | OTW | CAL/OTW |
| CC | GO:0005952 | cAMP-dependent protein kinase complex | 24 | 13 | 3.04 | 1.30E-06 | CAL | OTW | CAL/OTW |
| CC | GO:0005868 | cytoplasmic dynein complex | 36 | 16 | 4.56 | 2.40E-06 | CAL | OTW | CAL/OTW |
| CC | GO:0030687 | preribosome, large subunit precursor | 29 | 14 | 3.67 | 3.10E-06 | CAL | OTW | CAL/OTW |
| CC | GO:0000177 | cytoplasmic exosome (RNase complex) | 20 | 10 | 2.53 | 5.70E-05 | CAL | OTW | CAL/OTW |
| CC | GO:0034464 | BBSome | 14 | 8 | 1.77 | 9.60E-05 | CAL | OTW | CAL/OTW |
| CC | GO:0042555 | MCM complex | 14 | 8 | 1.77 | 9.60E-05 | CAL | OTW | CAL/OTW |
| CC | GO:0035869 | ciliary transition zone | 24 | 14 | 3.04 | 1.00E-04 | CAL | OTW | CAL/OTW |
| CC | GO:0036064 | ciliary basal body | 65 | 19 | 8.23 | 3.00E-04 | CAL | OTW | CAL/OTW |
| CC | GO:0000176 | nuclear exosome (RNase complex) | 24 | 10 | 3.04 | 0.00037 | CAL | OTW | CAL/OTW |
| CC | GO:0036038 | MKS complex | 13 | 7 | 1.65 | 0.00044 | CAL | OTW | CAL/OTW |
| CC | GO:0005615 | extracellular space | 95 | 24 | 12.03 | 0.00058 | CAL | OTW | CAL/OTW |
| CC | GO:0005858 | axonemal dynein complex | 18 | 8 | 2.28 | 0.00087 | CAL | OTW | CAL/OTW |
| CC | GO:0005856 | cytoskeleton | 528 | 106 | 66.88 | 0.00088 | CAL | OTW | CAL/OTW |
| MF | GO:0005524 | ATP binding | 1459 | 824 | 733.68 | 3.10E-07 | SHL | CAL | SHL/CAL |
| MF | GO:0008270 | zinc ion binding | 573 | 330 | 288.14 | 2.00E-04 | SHL | CAL | SHL/CAL |
| MF | GO:0005515 | protein binding | 3394 | 1805 | 1706.73 | 0.00042 | SHL | CAL | SHL/CAL |
| MF | GO:0003723 | RNA binding | 900 | 492 | 452.58 | 0.0033 | SHL | CAL | SHL/CAL |
| MF | GO:0003677 | DNA binding | 895 | 491 | 450.06 | 0.00458 | SHL | CAL | SHL/CAL |
| MF | GO:0016887 | ATP hydrolysis activity | 365 | 208 | 183.55 | 0.00545 | SHL | CAL | SHL/CAL |
| MF | GO:0003700 | DNA-binding transcription factor activit... | 261 | 153 | 131.25 | 0.00788 | SHL | CAL | SHL/CAL |
| MF | GO:0003676 | nucleic acid binding | 2595 | 1220 | 957.62 | 3.90E-19 | SHL | OTW | SHL/OTW |
| MF | GO:0008270 | zinc ion binding | 649 | 330 | 239.5 | 8.80E-14 | SHL | OTW | SHL/OTW |
| MF | GO:0003723 | RNA binding | 971 | 468 | 358.32 | 9.50E-13 | SHL | OTW | SHL/OTW |
| MF | GO:0046872 | metal ion binding | 1594 | 795 | 588.23 | 1.50E-09 | SHL | OTW | SHL/OTW |
| MF | GO:0016887 | ATP hydrolysis activity | 408 | 208 | 150.56 | 2.90E-09 | SHL | OTW | SHL/OTW |
| MF | GO:0005509 | calcium ion binding | 371 | 191 | 136.91 | 4.80E-09 | SHL | OTW | SHL/OTW |
| MF | GO:0005525 | GTP binding | 305 | 154 | 112.55 | 6.90E-07 | SHL | OTW | SHL/OTW |
| MF | GO:0016491 | oxidoreductase activity | 1280 | 565 | 472.35 | 3.80E-06 | SHL | OTW | SHL/OTW |
| MF | GO:0004672 | protein kinase activity | 640 | 312 | 236.18 | 6.80E-06 | SHL | OTW | SHL/OTW |
| MF | GO:0003924 | GTPase activity | 259 | 130 | 95.58 | 7.30E-06 | SHL | OTW | SHL/OTW |
| MF | GO:0004674 | protein serine/threonine kinase activity | 325 | 160 | 119.93 | 1.50E-05 | SHL | OTW | SHL/OTW |
| MF | GO:0004601 | peroxidase activity | 206 | 106 | 76.02 | 1.90E-05 | SHL | OTW | SHL/OTW |
| MF | GO:0003700 | DNA-binding transcription factor activit... | 292 | 152 | 107.76 | 6.40E-05 | SHL | OTW | SHL/OTW |
| MF | GO:0003729 | mRNA binding | 167 | 85 | 61.63 | 0.00014 | SHL | OTW | SHL/OTW |
| MF | GO:0061630 | ubiquitin protein ligase activity | 150 | 77 | 55.35 | 2.00E-04 | SHL | OTW | SHL/OTW |
| MF | GO:0000981 | DNA-binding transcription factor activit... | 150 | 77 | 55.35 | 2.00E-04 | SHL | OTW | SHL/OTW |
| MF | GO:0004843 | cysteine-type deubiquitinase activity | 123 | 65 | 45.39 | 0.00021 | SHL | OTW | SHL/OTW |
| MF | GO:0008017 | microtubule binding | 153 | 78 | 56.46 | 0.00025 | SHL | OTW | SHL/OTW |
| MF | GO:0051082 | unfolded protein binding | 132 | 68 | 48.71 | 0.00041 | SHL | OTW | SHL/OTW |
| MF | GO:0005524 | ATP binding | 1443 | 808 | 722.04 | 9.30E-07 | OTW | CAL | OTW/CAL |
| MF | GO:0005515 | protein binding | 3424 | 1814 | 1713.28 | 3.60E-05 | OTW | CAL | OTW/CAL |
| MF | GO:0008270 | zinc ion binding | 562 | 319 | 281.21 | 0.00064 | OTW | CAL | OTW/CAL |
| MF | GO:0003677 | DNA binding | 891 | 485 | 445.83 | 0.00202 | OTW | CAL | OTW/CAL |
| MF | GO:0003677 | DNA binding | 978 | 393 | 308.17 | 4.30E-21 | OTW | SHL | OTW/SHL |
| MF | GO:0004674 | protein serine/threonine kinase activity | 325 | 165 | 102.41 | 6.40E-12 | OTW | SHL | OTW/SHL |
| MF | GO:0004672 | protein kinase activity | 640 | 321 | 201.66 | 1.50E-11 | OTW | SHL | OTW/SHL |
| MF | GO:0003735 | structural constituent of ribosome | 267 | 134 | 84.13 | 1.20E-10 | OTW | SHL | OTW/SHL |
| MF | GO:0005524 | ATP binding | 1632 | 620 | 514.24 | 2.00E-09 | OTW | SHL | OTW/SHL |
| MF | GO:0022857 | transmembrane transporter activity | 840 | 317 | 264.68 | 4.90E-08 | OTW | SHL | OTW/SHL |
| MF | GO:0003755 | peptidyl-prolyl cis-trans isomerase acti... | 121 | 61 | 38.13 | 1.10E-05 | OTW | SHL | OTW/SHL |
| MF | GO:0008146 | sulfotransferase activity | 138 | 66 | 43.48 | 1.50E-05 | OTW | SHL | OTW/SHL |
| MF | GO:0031267 | small GTPase binding | 100 | 52 | 31.51 | 1.60E-05 | OTW | SHL | OTW/SHL |
| MF | GO:0005096 | GTPase activator activity | 99 | 51 | 31.19 | 2.60E-05 | OTW | SHL | OTW/SHL |
| MF | GO:0016757 | glycosyltransferase activity | 309 | 130 | 97.37 | 3.60E-05 | OTW | SHL | OTW/SHL |
| MF | GO:0046982 | protein heterodimerization activity | 95 | 49 | 29.93 | 3.60E-05 | OTW | SHL | OTW/SHL |
| MF | GO:0004222 | metalloendopeptidase activity | 99 | 50 | 31.19 | 6.20E-05 | OTW | SHL | OTW/SHL |
| MF | GO:0003713 | transcription coactivator activity | 64 | 35 | 20.17 | 1.00E-04 | OTW | SHL | OTW/SHL |
| MF | GO:0003682 | chromatin binding | 108 | 60 | 34.03 | 0.00017 | OTW | SHL | OTW/SHL |
| MF | GO:0051287 | NAD binding | 82 | 43 | 25.84 | 0.00032 | OTW | SHL | OTW/SHL |
| MF | GO:0016787 | hydrolase activity | 2321 | 650 | 731.34 | 0.00049 | OTW | SHL | OTW/SHL |
| MF | GO:0005085 | guanyl-nucleotide exchange factor activi... | 66 | 34 | 20.8 | 0.00056 | OTW | SHL | OTW/SHL |
| MF | GO:0003724 | RNA helicase activity | 66 | 34 | 20.8 | 0.00056 | OTW | SHL | OTW/SHL |
| MF | GO:0008146 | sulfotransferase activity | 141 | 73 | 17.55 | 2.50E-21 | CAL | SHL | CAL/SHL |
| MF | GO:0046982 | protein heterodimerization activity | 94 | 48 | 11.7 | 9.40E-20 | CAL | SHL | CAL/SHL |
| MF | GO:0045505 | dynein intermediate chain binding | 58 | 34 | 7.22 | 7.80E-17 | CAL | SHL | CAL/SHL |
| MF | GO:0030527 | structural constituent of chromatin | 71 | 36 | 8.84 | 5.00E-15 | CAL | SHL | CAL/SHL |
| MF | GO:0003677 | DNA binding | 895 | 203 | 111.38 | 2.90E-14 | CAL | SHL | CAL/SHL |
| MF | GO:0051959 | dynein light intermediate chain binding | 30 | 20 | 3.73 | 6.20E-12 | CAL | SHL | CAL/SHL |
| MF | GO:0004364 | glutathione transferase activity | 42 | 23 | 5.23 | 5.60E-11 | CAL | SHL | CAL/SHL |
| MF | GO:0046921 | alpha-(1->6)-fucosyltransferase activity | 33 | 20 | 4.11 | 8.20E-11 | CAL | SHL | CAL/SHL |
| MF | GO:0010181 | FMN binding | 43 | 23 | 5.35 | 1.10E-10 | CAL | SHL | CAL/SHL |
| MF | GO:0003676 | nucleic acid binding | 2466 | 425 | 306.9 | 1.50E-10 | CAL | SHL | CAL/SHL |
| MF | GO:0017108 | 5'-flap endonuclease activity | 34 | 20 | 4.23 | 1.70E-10 | CAL | SHL | CAL/SHL |
| MF | GO:0004795 | threonine synthase activity | 17 | 12 | 2.12 | 4.50E-08 | CAL | SHL | CAL/SHL |
| MF | GO:0051879 | Hsp90 protein binding | 33 | 17 | 4.11 | 6.20E-08 | CAL | SHL | CAL/SHL |
| MF | GO:0015179 | L-amino acid transmembrane transporter a... | 34 | 17 | 4.23 | 1.10E-07 | CAL | SHL | CAL/SHL |
| MF | GO:0008484 | sulfuric ester hydrolase activity | 31 | 16 | 3.86 | 1.50E-07 | CAL | SHL | CAL/SHL |
| MF | GO:0004176 | ATP-dependent peptidase activity | 27 | 14 | 3.36 | 8.40E-07 | CAL | SHL | CAL/SHL |
| MF | GO:0042162 | telomeric DNA binding | 21 | 12 | 2.61 | 1.30E-06 | CAL | SHL | CAL/SHL |
| MF | GO:0003713 | transcription coactivator activity | 56 | 21 | 6.97 | 1.50E-06 | CAL | SHL | CAL/SHL |
| MF | GO:0045505 | dynein intermediate chain binding | 59 | 34 | 7.23 | 9.90E-17 | CAL | OTW | CAL/OTW |
| MF | GO:0030527 | structural constituent of chromatin | 72 | 35 | 8.82 | 3.80E-14 | CAL | OTW | CAL/OTW |
| MF | GO:0003677 | DNA binding | 891 | 209 | 109.13 | 1.40E-13 | CAL | OTW | CAL/OTW |
| MF | GO:0046983 | protein dimerization activity | 181 | 69 | 22.17 | 2.30E-13 | CAL | OTW | CAL/OTW |
| MF | GO:0003676 | nucleic acid binding | 2415 | 431 | 295.78 | 2.80E-13 | CAL | OTW | CAL/OTW |
| MF | GO:0043565 | sequence-specific DNA binding | 194 | 51 | 23.76 | 2.90E-13 | CAL | OTW | CAL/OTW |
| MF | GO:0051959 | dynein light intermediate chain binding | 32 | 20 | 3.92 | 2.70E-11 | CAL | OTW | CAL/OTW |
| MF | GO:0004364 | glutathione transferase activity | 42 | 23 | 5.14 | 4.00E-11 | CAL | OTW | CAL/OTW |
| MF | GO:0010181 | FMN binding | 43 | 23 | 5.27 | 7.60E-11 | CAL | OTW | CAL/OTW |
| MF | GO:0017108 | 5'-flap endonuclease activity | 34 | 20 | 4.16 | 1.30E-10 | CAL | OTW | CAL/OTW |
| MF | GO:0020037 | heme binding | 92 | 34 | 11.27 | 9.80E-10 | CAL | OTW | CAL/OTW |
| MF | GO:0046921 | alpha-(1->6)-fucosyltransferase activity | 38 | 20 | 4.65 | 1.90E-09 | CAL | OTW | CAL/OTW |
| MF | GO:0004795 | threonine synthase activity | 15 | 12 | 1.84 | 3.50E-09 | CAL | OTW | CAL/OTW |
| MF | GO:0008484 | sulfuric ester hydrolase activity | 27 | 16 | 3.31 | 8.30E-09 | CAL | OTW | CAL/OTW |
| MF | GO:0051879 | Hsp90 protein binding | 31 | 17 | 3.8 | 1.40E-08 | CAL | OTW | CAL/OTW |
| MF | GO:0046982 | protein heterodimerization activity | 97 | 32 | 11.88 | 7.10E-08 | CAL | OTW | CAL/OTW |
| MF | GO:0004497 | monooxygenase activity | 87 | 37 | 10.66 | 2.90E-07 | CAL | OTW | CAL/OTW |
| MF | GO:0004176 | ATP-dependent peptidase activity | 26 | 14 | 3.18 | 3.70E-07 | CAL | OTW | CAL/OTW |
